# WNK1 enforces macrophage lineage fidelity

**DOI:** 10.1101/2023.04.26.538482

**Authors:** Alissa J. Trzeciak, Waleska Saitz Rojas, Zong-Lin Liu, Adam S. Krebs, Zhaoquan Wang, Pedro H. V. Saavedra, Isabella C. Miranda, Allie Lipshutz, Jian Xie, Chou-Long Huang, Michael Overholtzer, Michael S. Glickman, Christopher N. Parkhurst, Thomas Vierbuchen, Christopher D. Lucas, Justin S. A. Perry

## Abstract

The appropriate development of macrophages, the body’s professional phagocyte, is essential for organismal development, especially in mammals. This dependence is exemplified by the observation that loss-of-function mutations in colony stimulating factor 1 receptor (CSF1R) results in multiple tissue abnormalities owing to an absence of macrophages. Despite this importance, little is known about the molecular and cell biological regulation of macrophage development. Here, we report the surprising finding that the chloride-sensing kinase With-no-lysine 1 (WNK1) is required for development of tissue-resident macrophages (TRMs). Myeloid-specific deletion of *Wnk1* resulted in a dramatic loss of TRMs, disrupted organ development, systemic neutrophilia, and mortality between 3 and 4 weeks of age. Strikingly, we found that myeloid progenitors or precursors lacking WNK1 not only failed to differentiate into macrophages, but instead differentiated into neutrophils. Mechanistically, the cognate CSF1R cytokine macrophage-colony stimulating factor (M-CSF) stimulates macropinocytosis by both mouse and human myeloid progenitors and precursor cells. Macropinocytosis, in turn, induces chloride flux and WNK1 phosphorylation. Importantly, blocking macropinocytosis, perturbing chloride flux during macropinocytosis, and inhibiting WNK1 chloride-sensing activity each skewed myeloid progenitor differentiation from macrophages into neutrophils. Thus, we have elucidated a role for WNK1 during macropinocytosis and discovered a novel function of macropinocytosis in myeloid progenitors and precursor cells to ensure macrophage lineage fidelity.

**Highlights:**

- Myeloid-specific WNK1 loss causes failed macrophage development and premature death
- M-CSF-stimulated myeloid progenitors and precursors become neutrophils instead of macrophages
- M-CSF induces macropinocytosis by myeloid progenitors, which depends on WNK1
- Macropinocytosis enforces macrophage lineage commitment

## Introduction

Macrophages are innate immune cells that play an indispensable role as the body’s professional phagocyte that facilitate appropriate multicellular organismal development and maintain tissue homeostasis throughout life (Morioka et al., 2019; Wynn et al., 2013; Zago et al., 2021). As long-lived resident cells present in all tissues, tissue-resident macrophages (TRMs) are often the first line of defense against pathogens while simultaneously coordinating recruitment of effector cells that mediate the resolution of infection, dampening of associated inflammation, and subsequent tissue repair (Bosurgi et al., 2017; Ginhoux and Guilliams, 2016; Okabe and Medzhitov, 2016; Scott et al., 2014). Perhaps most importantly, TRMs are also responsible for the clearance of apoptotic cells, termed efferocytosis, a process that is critical during embryonic development and remains essential throughout life (Boada-Romero et al., 2020; Elliott and Ravichandran, 2016; Moore and Tabas, 2011; Trzeciak et al., 2021). The significance of TRM development and function is exemplified by the identification and investigation of the colony-stimulating factor 1 receptor (CSF1R; encoded by c*-fms* proto-oncogene) (Sherr et al., 1985). CSF1R is a receptor tyrosine kinase that signals in response to stimulation by macrophage colony-stimulating factor (M-CSF) or interleukin-34 (IL-34) (Easley-Neal et al., 2019; Lin et al., 2008; Ma et al., 2012; Stanley and Heard, 1977). Furthermore, CSF1R is expressed as early as embryonic day (E)6.5 in erythromyeloid progenitors and remains robustly expressed in multipotent progenitor cells (MPPs), mononuclear progenitor cells, monocytes, and all TRM subsets (Dai et al., 2002; Regenstreif and Rossant, 1989). Strikingly, mutation of CSF1, the cognate ligand to CSF1R, was shown to cause a near-complete loss of TRMs, which was found to account for the classic osteopetrotic (op/op) phenotype observed in mice, rats, and humans (Pixley and Stanley, 2004; Rojo et al., 2019; Wiktor-Jedrzejczak et al., 1990). Beyond studies of the CSF1–CSF1R axis, the bulk of work on TRM biology has understandably focused on the commitment and function of specific TRM subsets (Bleriot et al., 2020; Deczkowska et al., 2018; Olefsky and Glass, 2010). However, the molecular and cell biological factors that dictate macrophage lineage commitment remain largely unexplored.

With-no-lysine kinase 1 (WNK1) is a serine-threonine kinase that is important for cell volume regulation through its activity as a cytosolic chloride sensor (Yamada et al., 2016a; Yamada et al., 2016b). Recently, WNK1 and regulation of chloride flux were shown to be important for ‘healthy’ (e.g., non-inflammatory) efferocytosis *in vitro* (Perry et al., 2019) and for regulation of NLRP3 inflammasome activity in mature macrophages (Mayes-Hopfinger et al., 2021). Given these findings, we sought to explore if WNK1 was important for TRM homeostatic function using a strategy to conditionally delete *Wnk1* from TRMs prior to their development (at ~E6.5 during embryogenesis) (Deng et al., 2010). Unexpectedly, mice with *Csf1r* promoter-driven Cre− mediated deletion of *Wnk1* (WNK1-deficient) exhibited failure to thrive and were unable to live beyond four weeks of age. Unlike their littermate controls, WNK1-deficient mice broadly lacked TRMs across most organs assessed, displayed pathological tissue/organ development, and exhibited systemic neutrophil infiltration. Given 1) the striking similarity between the CSF1R mutant phenotype and WNK1-deficient mice, 2) that M-CSF/CSF1R signaling is essential for differentiation of myeloid progenitors and precursor cells into mature macrophages *in vitro* (Hamilton et al., 2014; Lacey et al., 2012), and 3) that M-CSF induces macropinocytosis in mature macrophages (Freeman et al., 2020; Racoosin and Swanson, 1989), we explored a novel hypothesis: differentiation of myeloid progenitors and precursor cells into macrophages depends on M-CSF-induced macropinocytosis, which itself depends on chloride flux and WNK1 activity. Here, we show that myeloid progenitors and precursor cells perform macropinocytosis in response to M-CSF, that macropinocytosis depends on WNK1, and that macropinocytosis is necessary for macrophage development. Strikingly, absence of WNK1, disruption of M-CSF-induced macropinocytosis, inhibition of WNK1 chloride-sensing activity, and perturbation of chloride flux during M-CSF-induced macropinocytosis, all prevented differentiation of macrophages from myeloid progenitors, instead driving progenitor differentiation into neutrophils. Thus, we have elucidated a role for WNK1 during macropinocytosis and discovered a novel function of macropinocytosis in myeloid progenitors and precursor cells to ensure macrophage lineage fidelity.

## Results

### WNK1 is required for tissue-resident macrophage development

To interrogate the importance of WNK1 for tissue-resident macrophage (TRM) function, we bred *Wnk1^fl/fl^* mice (Xie et al., 2009) to mice expressing Cre under the control of the colony stimulating factor 1 receptor (*Csf1r*) promoter. The resultant conditional loss of *Wnk1* in myeloid cells (*Csf1r^Cre+^;Wnk1^fl/fl^*, referred to as Cre+) resulted in significantly decreased fecundity (Figure S1) as well as dramatic premature death of all Cre+ mice between 3 and 4 weeks of age (**Figure 1A**). Additionally, Cre+ mice were significantly smaller than their *Csf1r^Cre–^;Wnk1^fl/fl^* (referred to as Cre−) control littermates (**Figure 1B**). Previous reports using the osteopetrotic (op) *Csf1*^op/op^ (Yoshida et al., 1990) and *Csf1r*-deficient (Dai et al., 2002) mice indicate that the CSF1-CSF1R pathway controls TRM development, including osteoclasts, which are required for bone development and remodeling (Jacome-Galarza et al., 2019). Interestingly, global CSF1R-deficient mice display lack of teeth, decreased body weight, and premature death (Dai et al., 2002). Similar to CSF1R-deficient mice, Cre+ mice lack tooth eruption (**Figure 1B**), suggesting that Cre+ mice also experience major defects in bone growth. Gross pathological examination of Cre+ mice showed that all major organs were smaller (**Figure 1B**) and histopathological analysis revealed evidence of malformation and inflammation (**Figure 1C** and S2B). Additionally, flow cytometry and immunofluorescence analysis of immune cells within vital organs (lungs, liver, kidney, heart, and brain) and lymphoid tissues (spleen and bone marrow) revealed TRM deficiency in all tissues except the brain (**Figures 1D** and **1E**; Figure S2A and S2C-E). Strikingly, alveolar, Kupffer, red pulp, intraglomerular, F4/80+ bone marrow (BM), and cardiac macrophages were largely absent in Cre+ mice (**Figure 1D** and **1E**; Figure S2C and S2D). Loss of TRMs in Cre+ mice was accompanied by a striking increase in Ly6G+ neutrophil infiltration across all tissues assessed (**Figure 1E** and **1F**; Figure S2D and S2E) as well as an increase in circulating neutrophils (Figure S2E and Data S1). Ly6C+ monocyte frequencies were either unaffected or modestly increased across tissues analyzed via flow cytometry and in complete blood count analysis (**Figure 1F** and S2E; Data S1). Subsequent analysis of monocyte subsets revealed differences in CX_3_CR1+ monocytes, including decreases in frequency in the spleen, liver, and lung (Figure S2F) suggesting that WNK1 may be required for the development of monocytes via a specific precursor (e.g., via GMP, but not MDP (Yáñez et al., 2017)). Additionally, we observed modest changes in the lymphoid compartment of some tissues (Figure S3A-C). The majority of CD4+ and CD8+ T cell numbers and frequencies remained similar across tissues analyzed, with only the spleen (increase in CD4+ T cells) and the lung (decrease in CD8+ T cells) showing significant changes (Figure S3B and S3C). We observed significantly decreased CD19+ B cell numbers and frequencies in nearly all tissues assessed (Figure S3A), however this defect is unlikely to explain our observed phenotype because mice lacking B cells develop normally (Kitamura et al., 1991). However, given the identification of a progenitor that has both B cell and myeloid potential and depends on CSF1R (Zriwil et al., 2016), it raises the possibility that B cell development via this progenitor may also depend on WNK1. Taken together, our findings suggest that mice bearing CSF1R^Cre^-specific deletion of WNK1 suffer perinatally-lethal defects in development due to failed tissue-resident macrophage (TRM) development.

**Figure 1:**
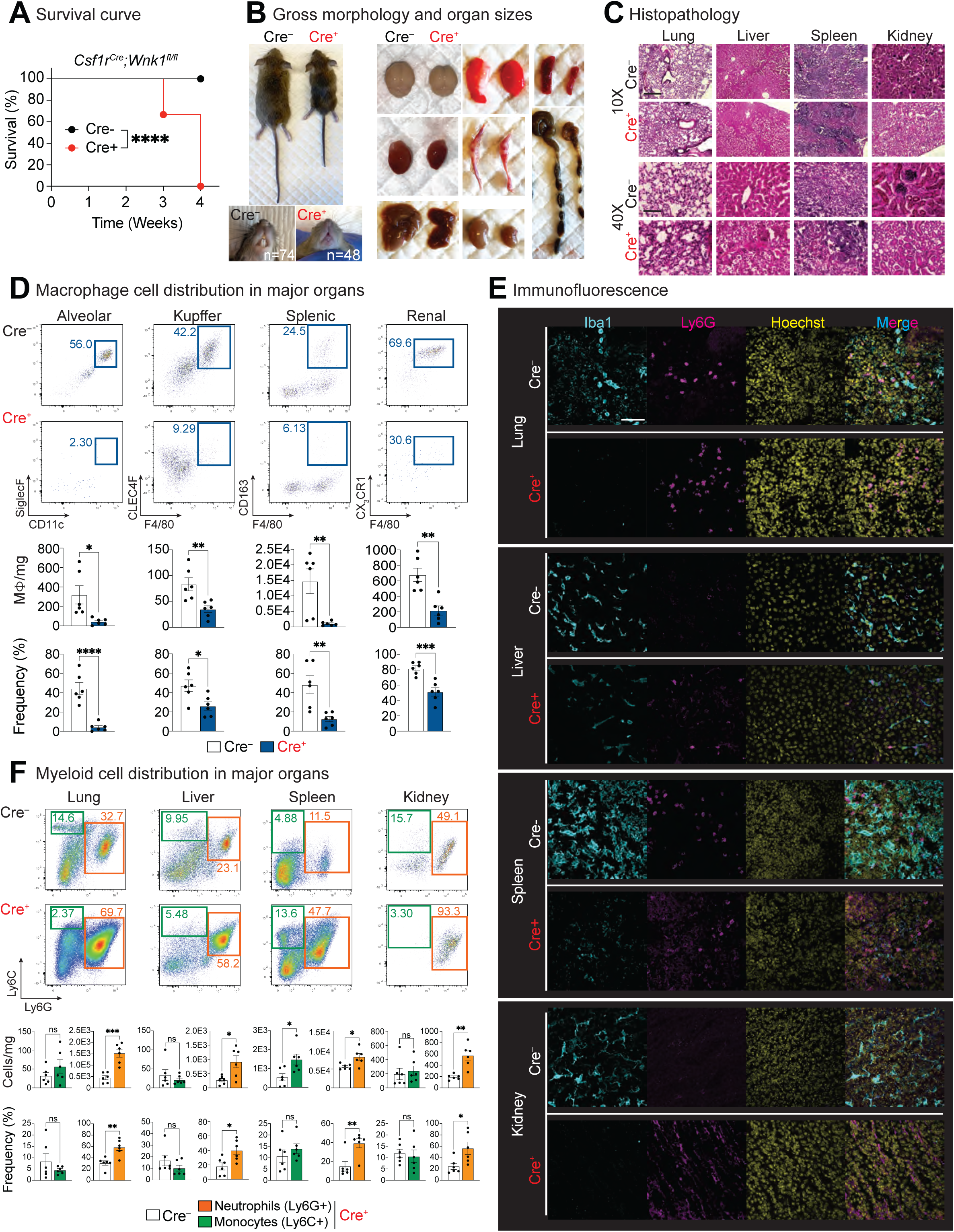
WNK1 is required for tissue-resident macrophage development. **(A)** <u>Myeloid cell-specific deletion of WNK1 results in early mortality</u>. Survival curve of *Csf1r^Cre+^;Wnk1^fl/fl^* (Cre+, red) vs. *Csf1^Cre–^;Wnk1^fl/fl^* (Cre−, black) mice. Litters were monitored from birth and sacrificed humanely when failure to thrive criteria were met. Data are from n=74 Cre− mice and n=48 Cre+ mice. Differences in survival were determined using the Mantel-Cox test. \*\*\*\**p* < .0001. See also Figure S1. **(B)** <u>Analysis of gross morphology and organ size reveals severe under-development</u>. (Left) Representative images of three- to four-week-old Cre− and Cre+ mice body (top) and teeth (bottom). Images are representative of n=74 Cre− mice and n=48 Cre+ mice. (Right) Representative images of organs from Cre− and Cre+ mice. Shown are (clockwise from top left) brain, lungs, spleen, colon, kidney, liver, heart, bones. Images are representative of organ analysis from six Cre− mice and six Cre+ mice. **(C)** <u>Histological analysis reveals disruption of tissue organization and inflammatory immune cell</u> <u>infiltration</u>. Hematoxylin & eosin (H&E) staining of lung, liver, spleen, and kidney. 10X scale bar, 300μm, 40X scale bar, 50 μm. Data are representative of four independent experiments. See also Figure S2B for analysis of additional tissues. **(D)** <u>Flow cytometric analysis of macrophage subsets in major organs</u>. Flow cytometry analysis of CD45+ tissue-resident macrophages in Cre+ (n=6) and Cre− (n=6) three-to-four-week-old mice. Subsets were first gated on live CD45+ CD11c-CD11b+ (except for lung macrophages; see Figure S2A for representative staining). Tissue-resident macrophages were gated as follows: alveolar macrophage (CD11c^high^ CD11b^low^ SiglecF+), Kupffer cells (Ly6C− Ly6G− CLEC4F+ F4/80+), splenic red pulp macrophages (Ly6C− Ly6G− CD163+ F4/80+), and kidney macrophages (Ly6C− Ly6G− CX_3_CR1+ F4/80+). Images (top) and summary plots of absolute numbers per milligram of tissue and frequencies (bottom) are from four independent experiments. Data are shown as mean ± SEM. Statistical significance was determined via independent samples *t*-test. \**p* < .05, \*\**p* < .01, \*\*\**p* < .001, \*\*\*\**p* < .0001, ns = not significant. See also Figure S2C for analysis of additional tissues. **(E)** <u>Immunofluorescence analysis of macrophages and neutrophils in the lung, liver, spleen, and</u> <u>kidney</u>. Confocal microscopy analysis was performed of lung, liver, spleen, and kidney macrophages (Iba1, cyan) and neutrophils (Ly6G, magenta). Nuclei were labeled using Hoechst (yellow). Scale bar, 20μm. Images are representative of four independent experiments. See also Figure S2D for analysis of additional tissues. **(F)** <u>Flow cytometric analysis of neutrophils and monocytes in major organs</u>. Flow cytometry analysis of CD45+ neutrophils and monocytes in Cre+ (n=6) and Cre− (n=6) three-to-four-week-old mice. Subsets were first gated on live CD45+ CD11c− CD11b+ then positively gated for neutrophils (orange bars, Ly6C+ Ly6G+) and monocytes (green bars, Ly6C^high^ Ly6G-). Images (top) and summary plots of absolute numbers per milligram of tissue and frequencies (bottom) are from four independent experiments. Data are shown as mean ± SEM. Statistical significance was determined via independent samples *t*-test. \**p* < .05, \*\**p* < .01, \*\*\**p* < .001, ns = not significant. See also Figure S2E. Monocytes were also analyzed for CX_3_CR1 expression (Figure S2F).

### Mice born with myeloid cells lacking WNK1 present with evidence of emergency myelopoiesis

The continued absence of TRMs and concomitant abundance of neutrophils suggest that the early mortality in Cre+ mice is due, at least in part, to emergency myelopoiesis in the hematopoietic compartment, similar to the changes observed in sepsis. To address this, we characterized the hematopoietic stem cell (HSC) niche in the bone marrow of Cre+ and littermate control mice (Figure S4A). Possibly owing to the difference in bone size between Cre+ and Cre− littermates, the absolute cell count of stem and progenitor populations were consistently lower in Cre+ animals (Figure S4B-E and Data S1). Unexpectedly, Cre+ mice exhibited a significant increase in frequency of the HSC subpopulations assessed (c-Kit^+^ Sca-1^+^, c-Kit^+^ Sca-1^−^ Figure S4B). Upon analysis of HSCs using CD48 and CD150 (SLAMF1) as classifiers of HSC stage (Figure S4C), we observed decreased long-term HSC frequency (LT; cKit^+^ Sca-1^+^ CD48^−^ CD150^+^), decreased (albeit not significant) short-term HSC frequency (ST; cKit^+^ Sca-1^+^ CD48^−^ CD150^−^), but increased multipotent progenitor frequency (MPP; cKit^+^ Sca-1^+^ CD48^+^ CD150^−^; Figure S4C). MPP subsets can be delineated first on expression of Fms Related Receptor Tyrosine Kinase 3 (Flt3), CD150, and CD48 into early or late MPPs (Figure S4D), and then into MPP2, MPP3, and MPP4 subsets, which give rise to erythrocyte/ megakaryocyte, granulocyte/ monocyte/ macrophage, and lymphoid lineages, respectively (Pietras et al., 2015) (Figure S4E). First, we found that Cre+ MPPs were predominantly Flt3^low^ (‘early’), seemingly less capable of progressing to the Flt3^high^ (‘late’) state (Figure S4D). Second, we found significant differences in both MPP3 and MPP4 populations but not in MPP2s. Specifically, we observed a higher frequency of MPP3s and a lower frequency of MPP4s in Cre+ mice compared to littermate controls (Figure S4E). Thus, absence of WNK1 leads to disrupted myelopoiesis reminiscent of pathological emergency myelopoiesis.

### WNK1 absence skews myelopoiesis towards neutropoiesis in response to M-CSF

Tissue-resident macrophages (TRMs), to some extent or another, can be repopulated by monocytes either endogenously or via adoptive transfer into neonatal mice (Jacome-Galarza et al., 2019). Despite having relatively normal (albeit phenotypically different) monocyte numbers (**Figure 1F**, S2E, and S2F), tissues from Cre+ mice still lack TRMs. Although this suggests that WNK1 is functioning downstream of CSF1R signaling cell-intrinsically, our findings could also be explained by disrupted tissue environments that are not conducive to macrophage differentiation and maturation. To test this alternative hypothesis, we injected either 1) wildtype monocytes or 2) wildtype multipotent progenitors into Cre+ mice and littermate controls beginning at either postnatal day (P)2 or P5 (**Figure 2A**). Injection of congenically-labeled wildtype monocytes into Cre+ mice starting at P5 was unable to rescue failure to thrive observed in Cre+ mice (**Figure 2B**, compare orange to blue). However, injection of congenically-labeled wildtype monocytes or multipotent progenitors into Cre+ mice beginning at P2 reversed failure to thrive in Cre+ mice (**Figure 2B** and **2C**, compare purple to blue). Consistent with these overt observations, flow cytometric analysis of key organs and tissues revealed significant rescue of the tissue macrophage pool by congenically-labeled wildtype monocytes or multipotent progenitors (**Figure 2D** and **2E**; Figure S5A). We next queried if monocyte or multipotent progenitor transfer also affected the neutrophilia observed in Cre+ mice (**Figure 1E** and **1F**). Most tissues and organs analyzed still exhibited significant neutrophilia, including the lungs and liver (Figure S5B-D), suggesting that TRM absence, and not neutrophilia per se, is the primary contributor to the mortality observed in *Csf1r^Cre+^;Wnk1^fl/fl^*(Cre+) mice. Thus, myeloid progenitors and precursor cells, including monocytes, that lack WNK1 fail to develop into macrophages *in vivo* which is a key contributor to the mortality observed in Cre+ mice.

**Figure 2.**
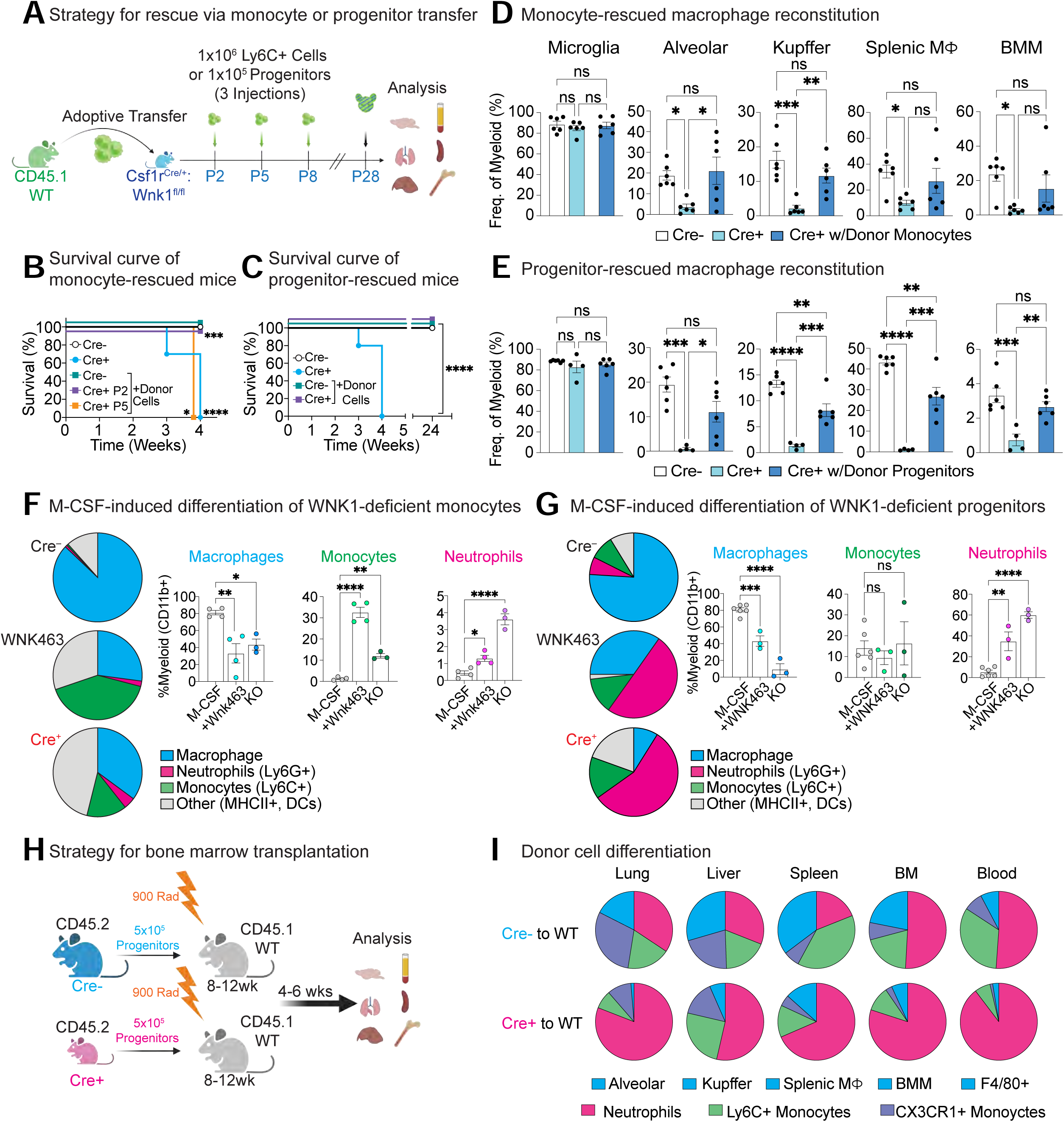
WNK1 absence skews myelopoiesis towards neutrophilia. **(A)** <u>Rescue of macrophage development via neonatal injection of bone marrow monocytes or</u> <u>myeloid progenitors</u>. Experimental strategy for bone marrow monocyte (**B, D**) or myeloid progenitor (**C, E**) adoptive transfer into *Csf1r^Cre+^;Wnk1^fl/fl^* (Cre+) or *Csf1^Cre–^;Wnk1^fl/fl^*(Cre−) neonatal mice. **(B, C)** <u>Analysis of survival in mice from experiments detailed in</u> Figure 2A. **(B)** Survival curve of Cre− mice (black), Cre+ mice (cyan), Cre− mice injected with wildtype CD45.1+ monocytes (green), and Cre+ mice injected with wildtype CD45.1+ monocytes beginning on perinatal day (P)2 (purple) or P5 (orange). Mice were monitored daily from birth and euthanized if IACUC-approved criteria were met. **(C)** Similar to **(B)**, except with myeloid progenitors injected at P2. Data are from two independent experiments with three mice per group. Differences in survival were determined using the Mantel-Cox test. \**p* < .05, \*\*\**p* < .001, \*\*\*\**p* < .0001. **(D, E)** <u>Analysis of macrophage reconstitution in mice from experiments highlighted in</u> Figure 2A. Summary plots of macrophage frequency as a fraction of CD45.1+ CD11b+ (except lung) F4/80+ cells in the brain, lungs, liver, spleen, and bone marrow of Cre− (white), Cre+ (light blue), and Cre+ mice injected with **(D)** wildtype CD45.1+ monocytes, or **(E)** wildtype CD45.1+ myeloid progenitors on P2 (dark blue). Data are from two independent experiments with two-to-three mice per group. Data are shown as mean ± SEM. Statistical significance was determined via one-way ANOVA. \**p* < .05, \*\**p* < .01, \*\*\**p* < .001, \*\*\*\**p* < .0001, ns = not significant. See also Figure S5 for analysis of neutrophil and monocyte subsets. **(F)** <u>Analysis of lineage frequency in M-CSF-treated mouse bone marrow Ly6C+Monoytes</u>. Bone marrow Ly6C+ monocytes were isolated from *Csf1r^Cre+^;Wnk1^fl/fl^*(Cre+), Cre+ pre-treated with 10μM WNK protein inhibitor, WNK463 (1h), or *Csf1^Cre–^;Wnk1^fl/fl^* (Cre−) mice and treated with M-CSF for 7d. Cultures were subsequently analyzed for the presence of macrophages, monocytes, neutrophils, and dendritic cells (DCs). Live cells were first gated as CD45+ CD11b+. (Left) Shown are fraction of the whole pie charts of neutrophils (Ly6G+ Ly6C+ F4/80−; magenta), monocytes (Ly6G− F4/80− Ly6C+; green), macrophages (Ly6C− Ly6G− F4/80+; blue), and other/DCs (F4/80-MHC Class II+; grey). (Right) Shown are summary graphs of the above identified populations. Data are representative of three independent experiments. Summary plots are shown as mean ± SD Statistical significance was assessed via independent samples *t*-test. \*\*\**p* < .001, ns = not significant. **(G)** <u>Analysis of lineage frequency in M-CSF-treated mouse bone marrow myeloid progenitors</u>. Similar to **(F)**, except bone marrow myeloid progenitors were isolated from *Csf1r^Cre+^;Wnk1^fl/fl^* (Cre+), Cre+ pre-treated with 10μM WNK protein inhibitor, WNK463 (1h), or *Csf1^Cre–^;Wnk1^fl/fl^*(Cre−) mice and treated with M-CSF for 7d. Cultures were subsequently analyzed for the presence of macrophages, monocytes, neutrophils, and dendritic cells (DCs). Live cells were first gated as CD45+ CD11b+. (Left) Shown are fraction of the whole pie charts of neutrophils (Ly6G+ Ly6C+ F4/80-; magenta), monocytes (Ly6G− F4/80− Ly6C+; green), macrophages (Ly6C− Ly6G− F4/80+; blue), and other/DCs (F4/80-MHC Class II+; grey). (Right) Shown are summary graphs of the above identified populations. Data are representative of three independent experiments. Summary plots are shown as mean ± SD Statistical significance was assessed via independent samples *t*-test. \*\*\**p* < .001, ns = not significant. **(H, I)** <u>Transplantation of Cre− or Cre+ myeloid progenitors into adult wildtype hosts</u>. **(H)** Schematic of experimental strategy. Myeloid progenitors from Cre− or Cre+ mice were transplanted into lethally irradiated (900rad) adult wildtype recipient mice, then analyzed 4-6 weeks post transfer. **(I)** Shown is the frequency of macrophages (blue), neutrophils (pink), Ly6C+ monocytes (green), and CX_3_CR1+ monocytes (purple) arising from Cre− (Cre− to WT) and Cre+ (Cre+ to WT) donor progenitors across tissues analyzed. Data are from two independent experiments with six (Cre+) or four (Cre−) mice per group. See also Figure S7 for statistical analyses of individual myeloid populations between Cre− and Cre+ conditions.

Macrophage colony-stimulating factor (M-CSF) is a cytokine critical for myeloid progenitor and precursor (i.e., monocyte) cell differentiation into macrophages instead of other immune cell lineages (Ushach and Zlotnik, 2016). Given our *in vivo* finding that Cre+ mice lack TRMs is reminiscent of CSF1R-deficient mice, we next tested if macrophage differentiation is affected in WNK1-deficient mice. To this end, we cultured bone marrow monocytes (**Figure 2F**, S6A and S6B) or bone marrow progenitors (**Figure 2G** and S6C) *in vitro* in the presence of M-CSF. Remarkably, cells from Cre+ mice were almost completely devoid of conventional mature macrophages (CD45+ CD11b+ F4/80+; **Figure 2G**, blue). Instead, Cre+ cells exhibited a predominantly neutrophilic phenotype (CD45+ CD11b+ Ly6C+ Ly6G+; **Figure 2G**, magenta). Contrarily, we observed no significant differences in monocytes (CD45+ CD11b+ Ly6G-Ly6C+, green) or dendritic cells (DCs; CD45+ F4/80-CD11c+ MHCII^high^, grey). Furthermore, this phenomenon could be induced in otherwise healthy cells, as addition of the pan-WNK inhibitor WNK463 (Yamada et al., 2016a; Yamada et al., 2016b; Zhang et al., 2016) to wildtype bone marrow progenitor cultures resulted in significantly decreased macrophage numbers and significantly increased neutrophil numbers (**Figure 2F and 2G**). We observed a similar finding when using bone marrow monocytes from Cre+ mice (**Figure 2F**). Specifically, we observed a significant reduction in conventional mature macrophages when Cre+ monocytes were treated with M-CSF (**Figure 2F**, blue). Unlike our findings with bone marrow progenitors, we observed a significant increase in monocytes when Cre+ monocytes were treated with M-CSF (**Figure 2F**, green). Surprisingly, we also observed a significant emergence of neutrophils in Cre+ monocytes treated with M-CSF (**Figure 2F**, pink), albeit at a much lower rate than observed when using bone marrow progenitors (**Figure 2G**, pink). This is likely a true conversion of monocytes into neutrophils given we started with a pure monocyte population devoid of bone marrow neutrophils. Additionally, bone marrow monocytes from wildtype mice treated with WNK463 recapitulated our findings with bone marrow monocytes from Cre+ mice (**Figure 2F**). Thus, our *in vitro* data suggest that the striking *in vivo* phenotype of TRM absence and neutrophilia is explained by myeloid progenitor and precursor cell dependence on WNK1 for appropriate lineage commitment in response to M-CSF.

Finally, we sought to directly test the hypothesis that Cre+ myeloid progenitors primarily differentiate into neutrophils instead of macrophages *in vivo*. We transferred progenitors from *Csf1r^Cre+^;Wnk1^fl/fl^* or littermate control mice into irradiated adult congenic C57BL/6J mice (**Figure 2H**). Recipient mice tissues were subsequently analyzed for the presence of donor macrophages, neutrophils, and monocyte subsets. Consistent with our *in vitro* findings, progenitors from *Csf1r^Cre+^;Wnk1^fl/fl^*mice failed to develop into TRMs, instead predominantly differentiating into neutrophils and monocytes (**Figure 2I** and S7A-D). These independent experiments demonstrate that, at least in early perinates, the tissues in *Csf1r^Cre+^;Wnk1^fl/fl^* mice (e.g., the bone marrow niche) remain capable of supporting macrophage development. Our findings suggest that the observed neutrophilia is not due to general inflammation but because of rerouted differentiation of myeloid progenitors from TRMs into neutrophils (as observed *in vitro*; **Figure 2F** and **2G**). Taken together, our data indicate that WNK1 is necessary for differentiation of myeloid progenitors and precursor cells into TRMs instead of neutrophils, and that the pathological consequence of myeloid-specific WNK1 absence is due to cell-intrinsic TRM development failure and not because of cell-extrinsic defects in the tissue microenvironment.

### WNK1 links M-CSF signaling and human macrophage differentiation

We next investigated whether the dependence on WNK1 for M-CSF-induced differentiation of myeloid progenitors into conventional macrophages is conserved in humans. To this end, we took two different approaches: M-CSF of either 1) peripheral blood CD14^high^ circulating monocytes or 2) human pluripotent stem cells in the presence of vehicle or the WNK functional inhibitor WNK463. First, peripheral blood CD14^high^ circulating monocytes cultured with M-CSF (**Figure 3A**) robustly produced morphologically and phenotypically conventional (CSF1R+) macrophages (**Figure 3B-D**). However, in the presence of WNK463, monocytes failed to differentiate into conventional macrophages (**Figure 3B-D** and S8A). It is worth noting that CD206 was significantly decreased (Figure S8A and S8B). This finding is similar to our observation with mouse bone marrow progenitors (Figure S6C) suggesting that expression of CD206, a protein associated with canonical anti-inflammatory responses, may require WNK1 (Perry et al., 2019).

**Figure 3.**
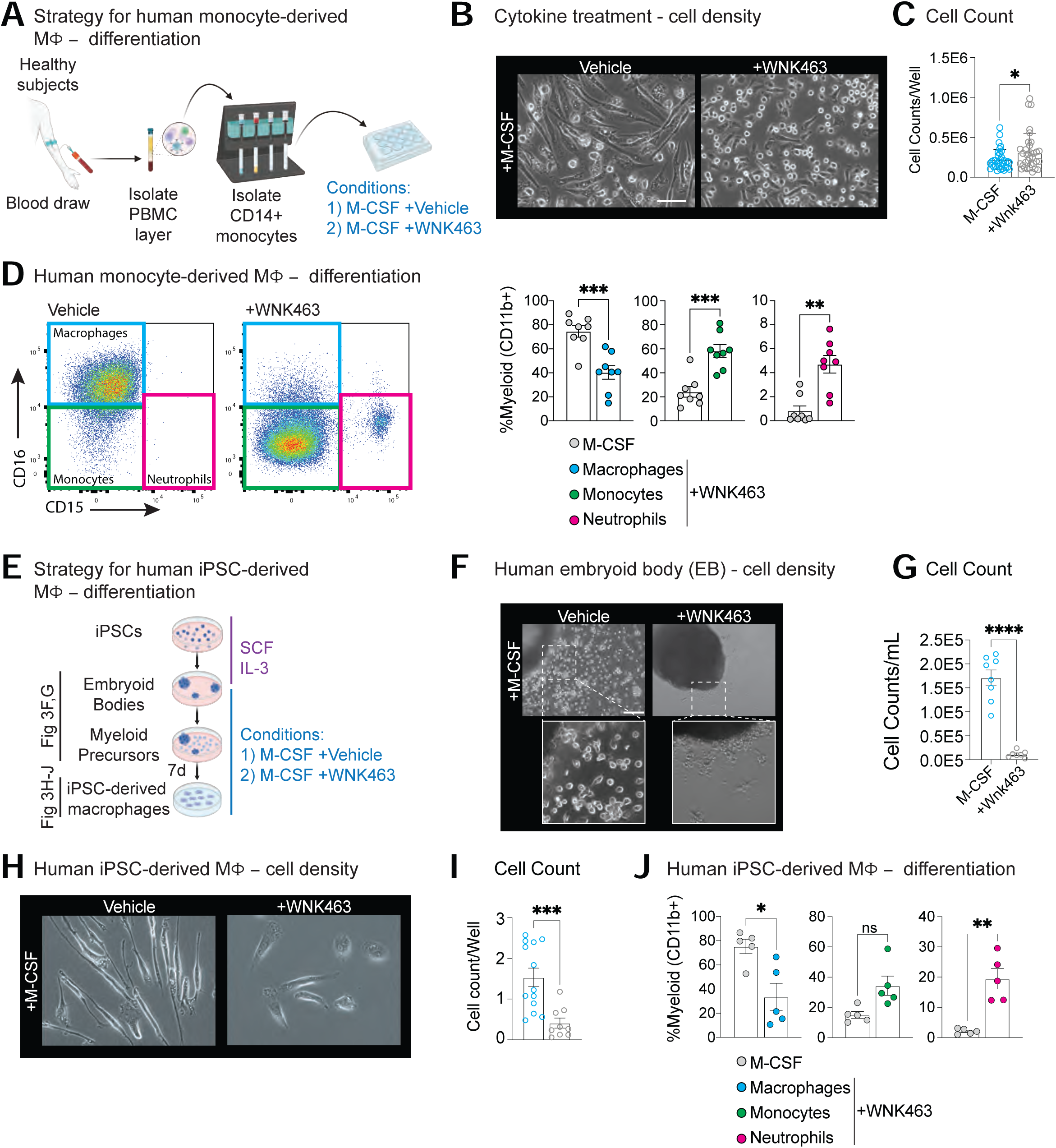
WNK1 supports differentiation of human myeloid progenitors and precursor cells into macrophages. **(A)** <u>Analysis of human blood monocyte-derived macrophage differentiation in response to M-CSF</u>. Shown is the experimental strategy used to test the effect of WNK protein function inhibition (via WNK463) on differentiation of human blood monocytes into macrophages. **(B-D)** <u>Perturbation of WNK function blocks maturation of human blood monocytes into mature</u> <u>macrophages</u>. Representative images **(B)**, quantitation of total cell numbers **(C)**, and flow cytometry analysis of myeloid subsets **(D)** from experiments performed as detailed in Figure 3A. In **(D)**, shown are representative flow cytometry (left) and summary (right) plots. Pooled data are from four independent experiments with two biological replicates (unique donors) per experiment. Data are shown as mean ± SEM. Statistical significance was determined via independent samples *t*-test. \**p* < .05, \*\**p* < .01, \*\*\**p* < .001. Scale bar, 150μm. See also Figure S8A for flow cytometry characterization of cell-surface proteins. **(E)** <u>Analysis of human pluripotent stem cell-derived macrophage differentiation in response to</u> <u>M-CSF</u>. Shown is the experimental strategy used to test the effect of WNK protein function inhibition (via WNK463) on differentiation of human pluripotent stem cells (PSCs) into macrophages. Also shown is the figure panel that corresponds to the state of differentiation being analyzed. **(F, G)** <u>Perturbation of WNK function blocks maturation of human embryoid bodies into myeloid</u> <u>progenitors</u>. Representative images **(F)** and quantitation **(G)** of myeloid progenitor numbers from experiments performed as detailed in Figure 3E. Pooled data are from four independent experiments with one-to-two biological replicates (unique donors) per experiment. Data are shown as mean ± SEM. Statistical significance was determined via independent samples *t*-test. \*\*\*\**p* < .0001. Scale bar, 150μm. **(H-J)** <u>Inhibition of WNK function prevents differentiation of human myeloid progenitors into</u> <u>mature macrophages</u>. Representative images **(H)**, quantitation of total cell numbers **(I)**, and **(J)** summary plots of myeloid subsets from experiments performed as detailed in Figure 3E. Pooled data are from four independent experiments with one-to-two biological replicates (unique donors) per experiment. Data are shown as mean ± SEM. Statistical significance was determined via independent samples *t*-test. \**p* < .05, \*\**p* < .01, \*\*\**p* < .001, ns = not significant. Scale bar, 20μm. See also Figure S8B for flow cytometry characterization of cell-surface proteins.

Next, we grew human pluripotent stem cell (hPSC)-derived embryoid bodies (EBs) in a defined media, which first differentiates EBs into myeloid precursors (via an M-CSF-dependent step), then terminally differentiates myeloid precursors into mature myeloid or granulocyte subsets (**Figure 3E**), including mature macrophages (Lim et al., 2013). Interestingly, we observed decreased M-CSF-dependent differentiation of EBs into myeloid progenitors when treated with WNK463 (**Figure 3F** and **3G**). As a complimentary approach, we first progressed EBs to myeloid progenitors prior to starting WNK463 treatment (**Figure 3E**). Myeloid progenitors were then treated with M-CSF in the presence of vehicle or WNK463. Similar to our results using mouse bone marrow progenitors, we found that treatment with WNK463 resulted in significantly fewer cells/lower cell density (**Figure 3H-J**) and reduced expression of conventional macrophage markers in response to M-CSF (Figure S8B). Collectively, our findings demonstrate that WNK1 dependence for M-CSF-induced myeloid progenitor and precursor cell differentiation into mature macrophages is conserved between mice and humans.

### Cellular morphology is altered in WNK1-deficient myeloid cells

Upon further examination of adherent Cre+ myeloid cells resulting from M-CSF treatment, we noted numerous conspicuous morphological differences. Specifically, M-CSF-treated Cre+ myeloid cells were significantly larger and presented a more circular morphology than M-CSF-treated Cre− myeloid cells, which featured a typical branching macrophage morphology (**Figure 4A** and **4B**). Strikingly, we also observed that adherent Cre+ myeloid cells resulting from M-CSF treatment contained significantly fewer intracellular vesicles than M-CSF-treated adherent Cre− myeloid cells (**Figure 4C**, left). Moreover, vesicles in adherent Cre+ myeloid cells were significantly smaller than vesicles in adherent Cre− myeloid cells (**Figure 4C**, right).

**Figure 4.**
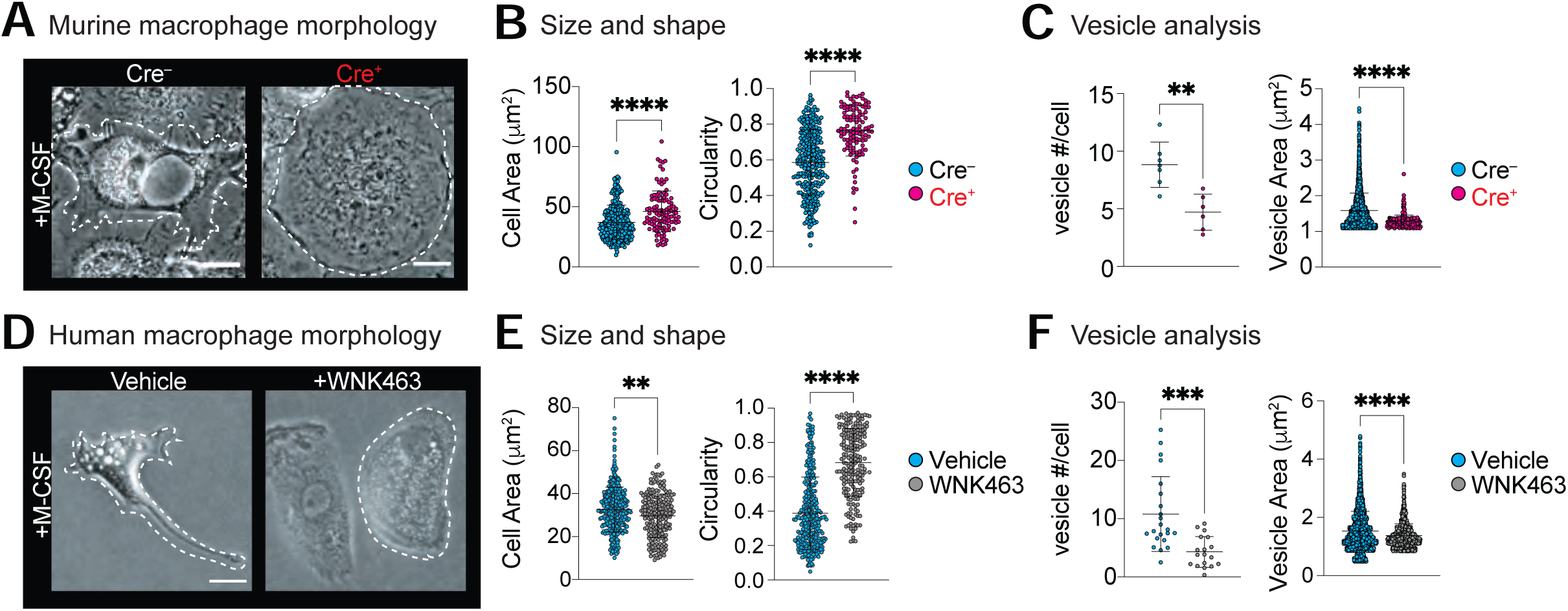
Cellular morphology is altered in WNK1-deficient myeloid cells. **(A, B)** <u>M-CSF-treated WNK1-deficient mouse myeloid cells exhibit altered cell morphology</u>. Representative images of cell morphology **(A)** and quantitation of cell size/shape **(B)** in myeloid cells from experiments performed as detailed in Figure 2F. Scale bars, 20μm. White dotted lines depict cell outlines. Area (left, per μm^2^) and circularity (right) were calculated after outlining cells as shown in Figure 4A. Images were blinded prior to analysis. Each data point represents a single cell. Data are pooled from three independent experiments. Data are shown as mean ± SD. Statistical significance was determined via independent samples *t*-test. \*\*\*\**p* < .0001. **(C)** <u>M-CSF-treated WNK1-deficient mouse myeloid cells contain fewer and smaller vesicles</u>. Analysis of number of vesicles per cell (left) and vesicle area (right) from experiments performed as detailed in Figure 4A. Each data point represents either a single cell (number of vesicles per cell) or a single vesicle (vesicle area). Data are pooled from three independent experiments. Data are shown as mean ± SD. Statistical significance was determined via independent samples *t*-test. \*\**p* < .01, \*\*\*\**p* < .0001. **(D, E)** <u>Human pluripotent stem cell (PSC)-derived myeloid cells exhibit altered cell morphology</u> <u>when WNK function is disrupted</u>. Representative images of cell morphology **(D)** and quantitation of cell size/shape **(E)** in myeloid cells from experiments performed as detailed in Fig. 3A. Scale bars, 20μm. White dotted lines depict cell outlines. Area (left, per μm^2^) and circularity (right) were calculated after outlining cells as shown in Figure 4D. Images were blinded prior to analysis. Each data point represents a single cell. Pooled data are from three independent experiments. Data are shown as mean ± SD. Statistical significance was determined via independent samples *t*-test. \*\**p* < .01, \*\*\*\**p* < .0001. **(F)** <u>Human PSC-derived myeloid cells with disrupted WNK function contain fewer and smaller</u> <u>vesicles</u>. Analysis of number of vesicles per cell (left) and vesicle area (right) from experiments performed as detailed in Figure 4D. Each data point represents either a single cell (number of vesicles per cell) or a single vesicle (vesicle area). Data are pooled from three independent experiments. Data are shown as mean ± SD. Statistical significance was determined via independent samples *t*-test. \*\*\**p* < .001, \*\*\*\**p* < .0001.

Given our findings with murine progenitors, we next investigated if human PSC-derived macrophages exhibited a similar phenotype. Indeed, macrophages derived from hPSCs using M-CSF in the presence of WNK463 had altered cell size and shape (**Figure 4D** and **4E**) and were nearly devoid of vesicles, with the present vesicles being significantly smaller (**Figure 4F**). These findings suggest that WNK1-deficient myeloid cells have disrupted vesicular trafficking in response to M-CSF which could arise, for instance, from altered uptake of extracellular content.

### M-CSF stimulates macropinocytosis in myeloid progenitors in a WNK1-dependent manner

WNK1 was recently shown to function as a cytosolic chloride-sensing kinase that responds rapidly to internalization of apoptotic cells (Perry et al., 2019). Specifically, efflux of cytosolic chloride induces WNK1 phosphorylation, which in turn signals to downstream kinases to initiate a corrective influx of extracellular chloride (Gagnon and Delpire, 2012). Given that phosphorylation of WNK1 leads to its activation and subsequent phosphorylation of effector kinases (Richardson et al., 2008; Vitari et al., 2006), we next assessed whether M-CSF induces WNK1 phosphorylation directly. Specifically, we isolated bone marrow progenitors from wildtype mice, stimulated progenitors with M-CSF, and measured WNK1 phosphorylation (**Figure 5A**). We also included WNK463 treatment, because WNK463 binds to and blocks a conserved catalytic site, which features both the chloride binding domain and activating phosphorylation sites, allowing us to also test the importance of chloride sensing by WNK1 (Yamada et al., 2016a; Yamada et al., 2016b; Zhang et al., 2016). Importantly, addition of M-CSF induced rapid (1h) WNK1 phosphorylation in myeloid progenitors (**Figure 5B**). Thus, WNK1, including its chloride sensing-dependent domain, is phosphorylated in response to M-CSF.

**Figure 5.**
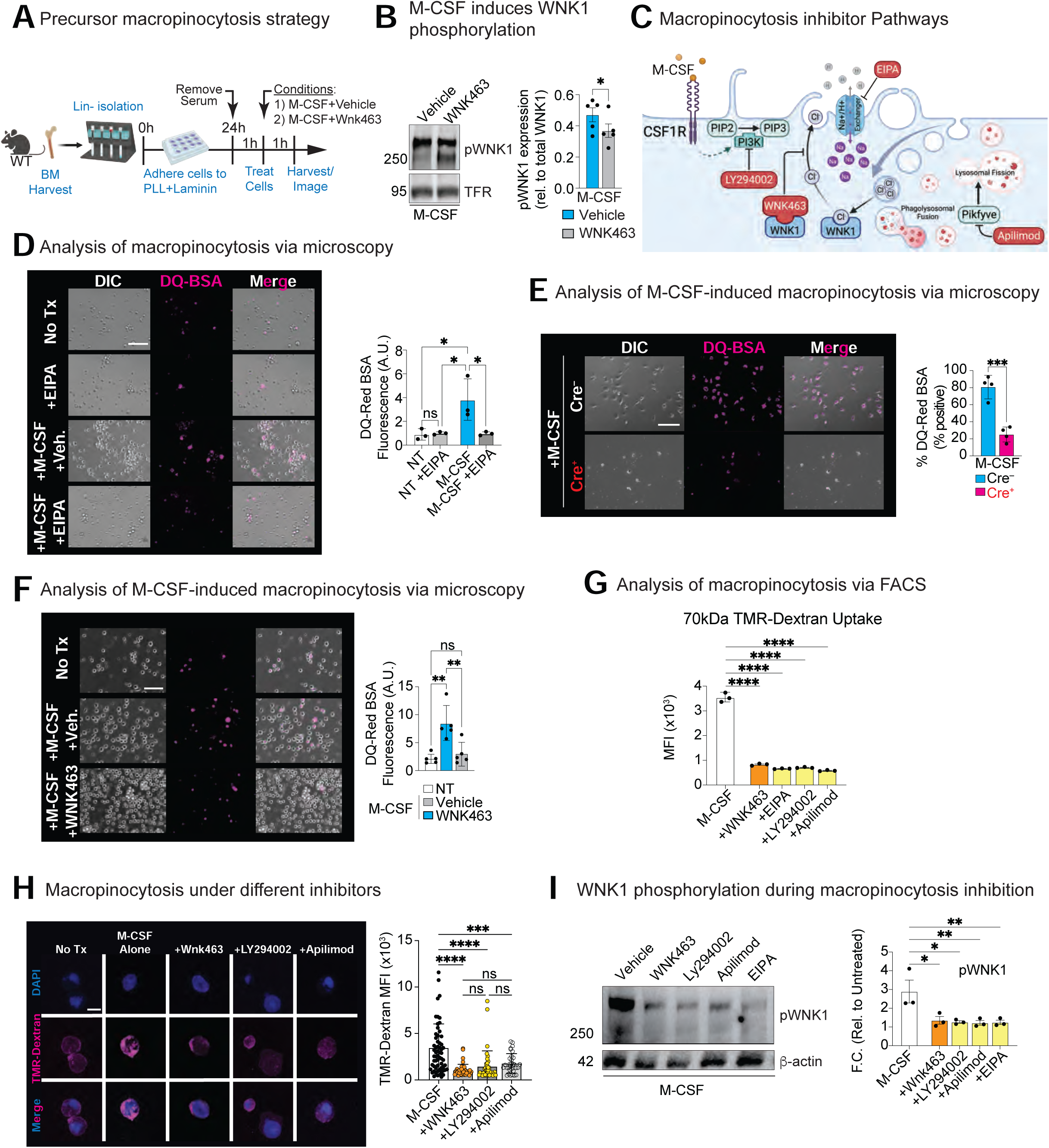
M-CSF stimulates macropinocytosis in myeloid progenitors in a WNK1-dependent manner. **(A)** <u>Strategy for analyzing M-CSF-induced macropinocytosis in bone marrow myeloid progenitors</u>. Shown is the experimental strategy used to determine if myeloid progenitors perform macropinocytosis in response to M-CSF and if WNK1 is required for this process. **(B)** <u>WNK1 catalytic site is phosphorylated in bone marrow myeloid progenitors in response to M-CSF</u>. Bone marrow myeloid progenitors were isolated and stimulated as in **(A)** in the presence of vehicle or WNK463, harvested after 1h, then assayed for WNK1 phosphorylation. Shown is a representative Phos-tag western blot (left) and quantitation of WNK1 phosphorylation (pWNK1; right). pWNK1 expression was calculated as a ratio of the single pWNK1 band to the non-phosphorylated WNK1 band (250kD), and then normalized to loading control (transferrin receptor; TFR). Data are from two independent experiments, with each experiment consisting of cells from at least two mice per condition (n=4 per condition). Data are shown as mean ± SEM. Statistical significance was determined via independent samples *t*-test. \**p* < .05. **(C)** <u>Methods for disrupting macropinocytosis</u>. Shown is a model of macropinocytosis with strategies used to disrupt this process. PIP2 - Phosphatidylinositol 4,5-bisphosphate. PIP3 - Phosphatidylinositol (3,4,5)-trisphosphate. PI3K - Phosphoinositide 3-kinase. PIKfyve - Phosphoinositide kinase, FYVE-type zinc finger containing. **(D)** <u>Bone marrow myeloid progenitors perform macropinocytosis in response to M-CSF</u>. Analysis of DQ-Red BSA internalization in bone marrow myeloid progenitors isolated and stimulated as in **(A)**. In conjunction with M-CSF, the putative macropinocytosis substrate DQ-Red bovine serum albumin (BSA) was added to cells and incubated for 1h in the presence of either vehicle or the macropinocytosis inhibitor 5-[N-ethyl-N-isopropyl] amiloride (EIPA). Representative microscopy images (left) and quantitation (fluorescence intensity in arbitrary units (A.U.); right) of DQ-Red BSA puncta are shown. Data is representative of two independent experiments with three biological replicates per experiment (n=6 per condition). Data are shown as mean ± SD. Statistical significance was determined via one-way ANOVA. \**p* < .05, ns = not significant. Scale bars, 50mm. **(E)** <u>WNK1 is required for M-CSF-induced macropinocytosis by bone marrow myeloid progenitors</u>. Bone marrow myeloid progenitors from *Csf1r^Cre+^;Wnk1^fl/fl^* (Cre+) or *Csf1^Cre–^;Wnk1^fl/fl^* (Cre−) mice were isolated and stimulated with M-CSF and DQ-Red BSA as in **(D)**. Representative microscopy images (left) and flow cytometry quantitation (right) of macropinocytotic (DQ-Red BSA+) progenitors are shown. Data is from two independent experiments with two mice (biological replicates) per experiment (n=4 per condition). Statistical significance was determined via independent samples *t*-test. \*\*\**p* < .001. Scale bars, 50mm. **(F)** <u>WNK kinase activity is required for M-CSF-induced macropinocytosis by bone marrow myeloid</u> <u>progenitors</u>. Wildtype bone marrow myeloid progenitors were isolated and stimulated with M-CSF and DQ-Red BSA as in **(D)**, except in the presence of vehicle or WNK463. Representative microscopy images (left) and quantitation (fluorescence intensity in arbitrary units (A.U.); right) of DQ-Red BSA+ puncta are shown. Data is representative of two independent experiments with at least two-to-three mice (biological replicates) per experiment (n=5 per condition). Data are shown as mean ± SD. Statistical significance was determined via one-way ANOVA (Bottom). \*\**p* < .01, ns = not significant. Scale bars, 50mm. **(G)** <u>M-CSF stimulates internalization of 70kDa Dextran by bone marrow myeloid progenitors via</u> <u>WNK1-dependent macropinocytosis</u>. Experiments performed as in **(D)**, but instead of DQ-Red BSA, the putative macropinocytosis substrate 70kDa TMR-Dextran was added together with M-CSF for 1h. Additionally, the WNK kinase activity inhibitor WNK463 or the macropinocytosis inhibitors EIPA, LY294002, or Apilimod (see Figure 5C) were added together with M-CSF. Shown is the summary plot of flow cytometry quantification of TMR-Dextran+ geometric mean fluorescence intensity (MFI). Data are from three independent experiments. Data are shown as mean ± SD. Statistical significance was determined via one-way ANOVA. \*\*\*\**p* < .0001. **(H)** <u>Time-lapse confocal microscopy analysis of M-CSF-induced internalization of 70kDa Dextran</u> <u>by bone marrow myeloid progenitors</u>. Experiments performed as in **(G)** but analyzed using time-lapse confocal microscopy over a 30min period. Representative microscopy images (left) and quantitation (fluorescence intensity in arbitrary units (A.U.); right) of TMR-Dextran+ myeloid progenitors are shown. Each dot is an individual cell, pooled from three independent experiments. Data are shown as mean ± SEM. Statistical significance was determined via one-way ANOVA. \*\*\**p* < .001, \*\*\*\**p* < .0001, ns = not significant. Scale bars, 5mm. **(I)** <u>Macropinocytosis triggers phosphorylation of the WNK1 catalytic site in bone marrow myeloid</u> <u>progenitors</u>. Bone marrow myeloid progenitors were isolated and stimulated for 1h as in **(B)**, but in the presence of vehicle, WNK463, or denoted macropinocytosis inhibitor (see Figure 5C), then assayed for WNK1 phosphorylation. Shown is a representative Phos-tag western blot (left) and quantitation of WNK1 phosphorylation (pWNK1; right). pWNK1 expression was calculated as in **(B)**, presented as a fold change (F.C.) relative to untreated myeloid progenitors. Data are from three independent experiments, with each experiment consisting of cells from at least one mouse per condition. Data are shown as mean ± SEM. Statistical significance was determined via one-way ANOVA. \**p* < .05. \*\**p* < .01, ns = not significant.

Interestingly, stimulation of macrophages with M-CSF triggers macropinocytosis, or “cell drinking”, an engulfment process first observed by Warren H. Lewis in 1931, yet the physiologic functions of macropinocytosis are only recently starting to emerge (Canton, 2018; King and Kay, 2019; Schmid et al., 2014; Swanson and King, 2019). Tissue-resident macrophages (TRMs) use M-CSF-induced macropinocytosis as a means of sensing their surroundings, but research on this type of uptake is limited to terminally-differentiated macrophages (Canton, 2018; Freeman et al., 2020; Mendel et al., 2022). Despite the known M-CSF dependence for macrophage differentiation, there is no evidence that myeloid progenitors or precursor cells perform macropinocytosis in response to M-CSF or that macropinocytosis is required for macrophage lineage commitment. Indeed, our observation that WNK1-deficient myeloid cells lack vesicles in response to M-CSF (**Figure 4**) suggests that 1) progenitors and precursor cells perform macropinocytosis and 2) progenitor and precursor cell macropinocytosis is disrupted in the absence of WNK1. To directly test this, we performed multiple complimentary experiments involving two separate macropinocytosis substrates (DQ-BSA and 70kDa Dextran) together with independent inhibitors of macropinocytosis and approaches of targeting WNK1 (**Figure 5C**).

First, to determine if bone marrow progenitors are capable of performing macropinocytosis in response to M-CSF, we cultured purified bone marrow progenitors with M-CSF and the macropinocytosis substrate DQ-Red BSA, a dye which fluoresces upon cleavage by cathepsin protease activity following uptake into the macropinosome (Marwaha and Sharma, 2017; Perry et al., 2019). Additionally, we included the macropinocytosis-selective inhibitor 5-[N-ethyl-N-isopropyl] amiloride (EIPA) (Jayashankar and Edinger, 2020; Kim et al., 2018; Koivusalo et al., 2010; Racoosin and Swanson, 1992). Strikingly, bone marrow progenitors performed significant macropinocytosis in response to M-CSF that was absent in unstimulated cells and completely abrogated by treatment with EIPA (**Figure 5D**).

Second, to target WNK1, we cultured bone marrow progenitors from *Csf1r^Cre+^;Wnk1^fl/fl^*(Cre+) mice or littermate controls (Cre−) with M-CSF and DQ-Red BSA. Stimulation with M-CSF induced robust uptake of DQ-Red BSA in Cre− bone marrow progenitors that was significantly decreased in Cre+ bone marrow progenitors (**Figure 5E**). To rule out potential confounds arising during stem cell development, we cultured bone marrow progenitors from wildtype mice with M-CSF and DQ-Red BSA in the presence of vehicle or WNK463. Stimulation with M-CSF induced robust uptake of DQ-Red BSA in vehicle-treated wildtype control myeloid progenitors that was significantly reduced when wildtype myeloid progenitors were treated with WNK463 (**Figure 5F**).

Third, because EIPA and WNK463 both target ion transport (and putatively lysosomal acidification), we tested M-CSF-induced macropinocytosis in bone marrow progenitors using the pH-insensitive macropinocytosis substrate 70kDa Dextran. Similar to our results with DQ-BSA, we found that M-CSF induces uptake of 70kDa Dextran by myeloid progenitors which is abrogated by WNK463 and EIPA (**Figure 5G**). Lastly, we targeted alternative pathways thought to support macropinocytosis (**Figure 5C**). Similar to our results with EIPA, inhibition of phosphatidylinositol 3-kinase (PI3K) using LY294002 or of Phosphoinositide Kinase, FYVE-Type Zinc Finger Containing (PIKfyve) using Apilimod significantly abrogated macropinocytotic uptake of 70kDa Dextran independently assessed using flow cytometry and high resolution time-lapse confocal microscopy (**Figure 5G** and **5H**). Taken together, our data suggest that M-CSF induces macropinocytosis in bone marrow progenitors and that WNK1 is required for M-CSF-induced macropinocytosis.

We next took advantage of our finding that M-CSF induces macropinocytosis in bone marrow progenitors in a WNK1-dependent manner to directly test if M-CSF-induced WNK1 phosphorylation requires macropinocytosis. As we observed previously (**Figure 5B**), incubation of progenitors with M-CSF induced robust WNK1 phosphorylation (**Figure 5I**, white bar). As hypothesized, inhibition of macropinocytosis using EIPA, LY294002, and Apilimod all significantly abrogated WNK1 phosphorylation (**Figure 5I**, yellow bars). Thus, our data indicate that M-CSF-stimulated macropinocytosis in bone marrow progenitors induces WNK1 signaling.

### Macropinocytosis is required for macrophage differentiation

Building on our finding that M-CSF triggers macropinocytosis by bone marrow progenitors in a WNK1-dependent manner, we sought to directly test the hypothesis that M-CSF-induced macropinocytosis is necessary for bone marrow progenitor differentiation into macrophages. Indeed, inhibition of macropinocytosis resulted in a significant decrease in myeloid progenitor differentiation into macrophages (F4/80+) and a concomitant increase in neutrophil generation (Ly6G+, **Figure 6A**). The skewed differentiation of bone marrow progenitors was more variable and less significant in EIPA-treated cells than in WNK463-treated cells (**Figure 6A**), which might relate to the potency of the inhibitor or suggest an additional function of WNK1 during myeloid differentiation.

**Figure 6.**
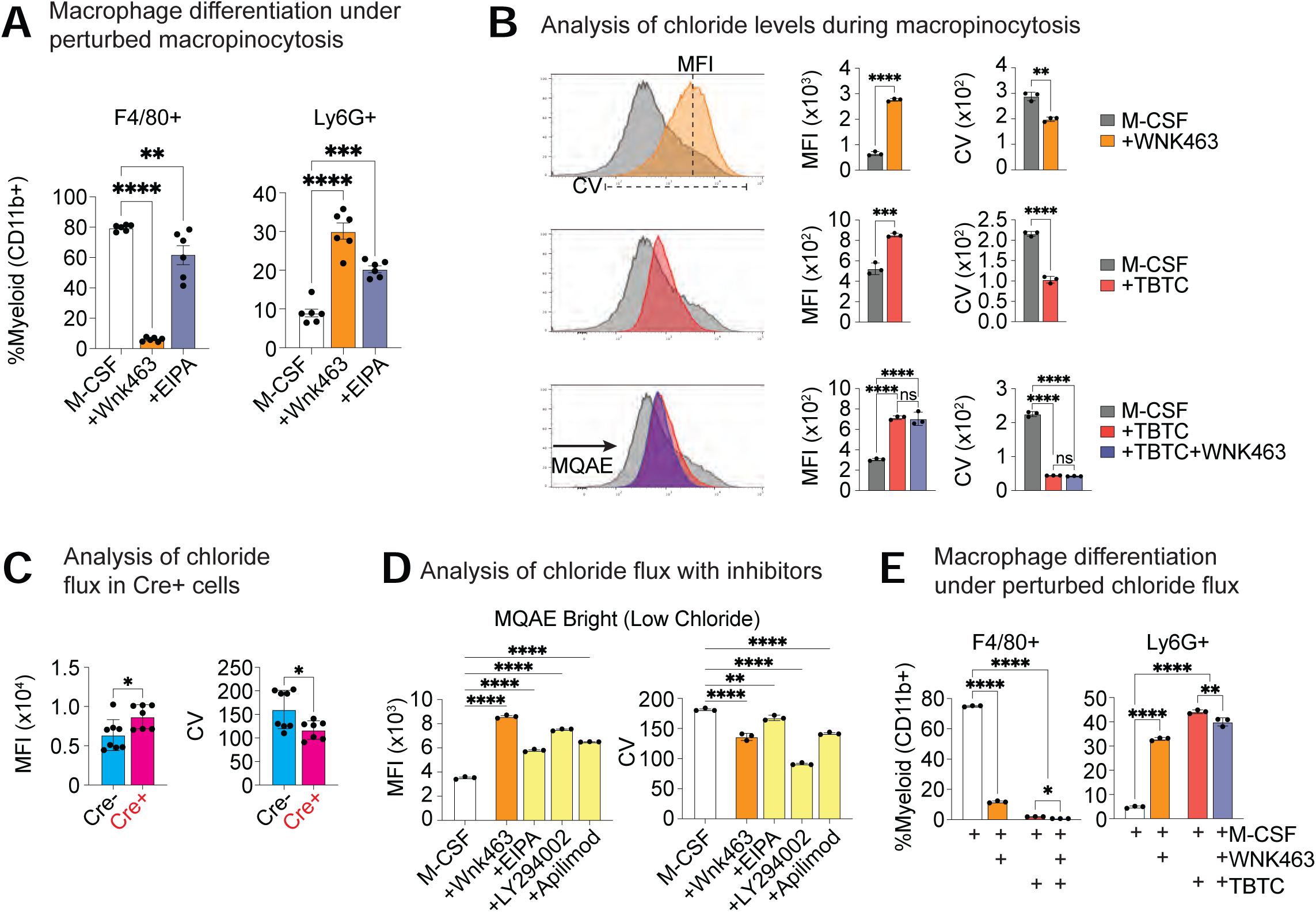
Macropinocytosis is required for macrophage differentiation. **(A)** <u>Perturbation of macropinocytosis skews M-CSF-stimulated bone marrow myeloid progenitor differentiation from macrophages to neutrophils</u>. Bone marrow myeloid progenitors were isolated and differentiated with M-CSF in the presence of vehicle, WNK463, or EIPA for 4d. Shown are summary plots of flow cytometry analysis for macrophages (F4/80+) and neutrophils (Ly6G+). Data are presented as a frequency of myeloid cells (CD11b+) from three independent experiments with two mice per experiment. Data are shown as mean ± SEM. Statistical significance was determined via one-way ANOVA. \*\**p* < .01, \*\*\**p* < .001, \*\*\*\**p* < .0001. **(B, C)** <u>M-CSF induces WNK1-dependent bidirectional chloride flux</u>. Either wildtype bone marrow myeloid progenitors **(B)** or bone marrow myeloid progenitors from *Csf1r^Cre+^;Wnk1^fl/fl^* (Cre+) or *Csf1^Cre–^;Wnk1^fl/fl^*(Cre−) mice **(C)** were isolated and labeled with the chloride indicator N-[ethoxycarbonylmethyl]-6-methoxy-quinolinium bromide (MQAE) prior to stimulation with M-CSF for 1h. MQAE fluorescence is inversely proportional to intracellular chloride level. For instance, a relative decrease in MQAE fluorescence corresponds to a relative increase in intracellular chloride (low chloride = MQAE bright). In **(B)**, myeloid progenitors were concomitantly treated with WNK463 (top) and/or the chloride ionophore tributyltin chloride (TBTC; middle and bottom). Shown are representative histograms (**B**, left) and summary plots (**B**, right and **C**) of the MQAE geometric mean fluorescence intensity (MFI) and the population distribution of fluorescence (coefficient of variation, CV). Data are representative of three independent experiments **(B)** or two pooled independent experiments (four replicates per experiment, **C)**. Data are shown as mean ± SD **(B)** or as mean ± SEM **(C)**. Statistical significance was determined via one-way ANOVA. \**p* < .05, \*\**p* < .01, \*\*\**p* < .001, \*\*\*\**p* < .0001, ns = not significant. **(D)** <u>Macropinocytosis induces bidirectional chloride flux</u>. Experiments were performed as in (**B**), but instead cultured in the presence of vehicle, WNK463, or denoted macropinocytosis inhibitor (see Figure 5C) during M-CSF stimulation. Shown are summary plots of the MQAE geometric mean fluorescence intensity (MFI) and the population distribution of fluorescence (coefficient of variation, CV). Data are from three independent experiments. Data are shown as mean ± SD. Statistical significance was determined via one-way ANOVA. \*\**p* < .01, \*\*\*\**p* < .0001. **(E)** <u>Perturbation of chloride flux skews M-CSF-stimulated bone marrow myeloid progenitor differentiation from macrophages to neutrophils</u>. Bone marrow myeloid progenitors were isolated and differentiated with M-CSF in the presence of vehicle, WNK463, TBTC, or WNK463+TBTC for 4d. Shown are summary plots of flow cytometry analysis for macrophages (F4/80+) and neutrophils (Ly6G+). Data are presented as a frequency of myeloid cells (CD11b+) from three independent experiments. data are shown as mean ± SD. Statistical significance was determined via one-way ANOVA. \**p* < .05, \*\**p* < .01, \*\*\**p* < .001, \*\*\*\**p* < .0001.

We next sought to further delineate how WNK1 is functioning downstream of M-CSF-induced macropinocytosis in bone marrow progenitors. Specifically, we sought to determine whether WNK1 functions in bone marrow progenitors to sense and modulate cytosolic chloride flux during M-CSF-induced macropinocytosis. To examine chloride flux in bone marrow progenitors, we used N-[ethoxycarbonylmethyl]-6-methoxy-quinolinium bromide (MQAE), a chemical that reversibly binds chloride, increasing in fluorescence intensity upon a relative decrease in cytosolic chloride (Koncz and Daugirdas, 1994; Verkman et al., 1989). Indeed, M-CSF-treated progenitors exhibited a broad distribution of MQAE fluorescence (Coefficient of Variation - CV, **Figure 6B**, top row) including a population of progenitors with high chloride (Geometric Mean Fluorescence Intensity - MFI, **Figure 6B**, top row). Interestingly, both inhibition of the conserved chloride-sensing domain of WNK proteins via WNK463 and absence of WNK1 significantly stifled both the broad distribution of MQAE fluorescence and the increased intracellular chloride levels observed in M-CSF-stimulated progenitors (**Figure 6B**, top row and **Figure 6C**). Given our finding that bone marrow progenitor macropinocytosis induces significant chloride influx, we next queried whether we could modulate chloride flux in actively macropinocytotic progenitors. To do so, we used the chloride ionophore tributyltin chloride (TBTC), which forces chloride into or out of the cell against its concentration gradient, disrupting typical chloride flux dynamics (Perry et al., 2019). Interestingly, TBTC significantly narrowed the range of MQAE fluorescence and decreased the cytosolic chloride levels in M-CSF-stimulated progenitors (**Figure 6B**, middle row). Furthermore, TBTC was able to force chloride into the cytosol of M-CSF-stimulated bone marrow progenitors despite having reduced macropinocytosis because of WNK463 treatment (**Figure 6B**, bottom row), confirming its utility as a tool to bypass M-CSF-induced, WNK1-regulated chloride flux. Our data suggest that M-CSF induces chloride flux, which subsequently requires WNK1 chloride-sensing activity. Finally, having established that M-CSF induces chloride flux, we tested whether macropinocytosis itself induces chloride flux in bone marrow progenitors when stimulated with M-CSF. Indeed, perturbing macropinocytosis via multiple methods (**Figure 5C**) by bone marrow progenitors significantly disrupted chloride flux similar to inhibition of WNK1 activity (**Figure 6D**). Thus, M-CSF-induced macropinocytosis by bone marrow progenitors triggers chloride flux, which is sensed and regulated by WNK1 via its chloride-sensing kinase activity.

Finally, we tested if disruption of appropriate chloride flux in response to M-CSF treatment affects macrophage differentiation. Strikingly, TBTC treatment of bone marrow progenitors stimulated with M-CSF completely blocked macrophage differentiation, instead resulting in increased differentiation of neutrophils (**Figure 6E**, compare red and white bars). Interestingly, treatment with TBTC, which bypasses WNK1 and induces chloride flux against the concentration gradient, was unable to overcome blockade of WNK kinase function (**Figure 6E**, compare dark blue and white bars). This data suggests that WNK1 chloride-sensing activity, and not chloride flux per se, supports differentiation of bone marrow progenitors into macrophages in response to M-CSF-induced macropinocytosis. Collectively, our results demonstrate that M-CSF-induced macropinocytosis by bone marrow progenitors is necessary for appropriate myeloid lineage commitment.

## Discussion

Tissue-resident macrophages (TRMs) are essential for organismal development and tissue homeostasis. TRM development is dependent on CSF1R signaling, but the cell biological and molecular factors that drive differentiation of myeloid progenitors into TRMs remain poorly understood. Here, we make several unexpected, novel observations relevant to myeloid progenitor and precursor cell and molecular biology, myeloid cell lineage commitment, and TRM macrophage development. First, across multiple lines of *in vivo* and *in vitro* evidence, we found that the serine-threonine kinase WNK1 is required for both mouse and human myeloid progenitor and precursor cell differentiation into macrophages. Second, M-CSF (via CSF1R) stimulates macropinocytosis, a phenomenon previously not observed in progenitors or precursor cells. Third, M-CSF-stimulated macropinocytosis induces WNK1 phosphorylation and activation, likely in response to chloride flux. Fourth, absence of WNK1, blockade of M-CSF-induced macropinocytosis, inhibition of chloride-sensing by WNK1, and altered chloride flux during M-CSF-induced macropinocytosis, all disrupt macrophage differentiation, instead driving differentiation towards the granulocyte lineage (e.g., neutrophils). Taken together, we have discovered a new phenomenon whereby macropinocytosis stimulates the differentiation of progenitors and precursor cells into a specific lineage and elucidated a molecular circuit by which this biological process ensures lineage fidelity.

Given the striking loss of TRMs across multiple tissues, we were initially surprised that monocyte numbers and frequencies were either unchanged or only modestly changed across tissues. However, our data is consistent with previous observations in op/op mice and mice lacking CSF1R (Cecchini et al., 1994; Dai et al., 2002; Lenzo et al., 2012; MacDonald et al., 2010). Importantly, we did observe changes in the frequency of specific monocyte subsets in some tissues. These findings, together with a previous report suggesting that phenotypically-distinct monocytes can develop via one of two independent progenitors (Yáñez et al., 2017), suggest that monocytes develop via a route independent of M-CSF/CSF1R (Greter et al., 2012; Hoeffel et al., 2015; Hoeffel and Ginhoux, 2018; MacDonald et al., 2010) but that these monocytes are subsequently unable to develop into TRMs in response to M-CSF. Future work will be required to determine the nature of this progenitor and what cell biological processes drive its development.

Both M-CSF and IL-34 signal through CSF1R, yet IL-34 selectively supports microglia and Langerhans cells development and maintenance (Greter et al., 2012; Wang et al., 2012). Despite increased neutrophilia in the brain, microglia numbers were relatively unchanged. Interestingly, IL-34 appears to not be required for embryonic development of microglia but instead for their maintenance. Indeed, IL-34-deficient mice exhibit reduced, not absent, adult microglia numbers (Greter et al., 2012; Wang et al., 2012). This raises several intriguing questions. For instance, are there two independent routes for generating microglia: one that is dependent on CSF1R and another that is not? Additionally, does IL-34 induce macropinocytosis in microglia progenitors and if so, is it qualitatively different than that in myeloid progenitors? Interestingly, a recent report found that IL-34 is capable of inducing macropinocytosis in mature macrophages *in vitro* but is inhibited by the presence of certain amino acids (Mendel et al., 2022), suggesting that the nutrient environment may also inform induction and reliance on macropinocytosis in progenitors. Finally, are the developing skin and CNS environments, both of which arise from the ectoderm, uniquely contributing factors that inform TRM differentiation in conjunction with IL-34? Combining our WNK1 floxed mice together with newly developed Langerhans cell- and microglia-specific Cre strains may provide a means for testing these intriguing hypotheses.

Unlike other lysosomal degradative processes (e.g., efferocytosis, receptor-mediated endocytosis, autophagy), the biological importance of macropinocytosis remains less understood. This is particularly surprising given that macropinocytosis is highly evolutionarily conserved and has been coopted by immune cells and cancer for unique functional purposes (King and Kay, 2019). To date, macropinocytosis has primarily been reported to be important for growth factor signaling in lower organisms (e.g., Dictyostelium), in differentiated mammalian cells (e.g., macrophages), and in oncogenically transformed cells (King and Kay, 2019; Swanson and King, 2019). However, an essential physiological role for macropinocytosis in metazoans has remained elusive. Here, we not only provide the first demonstration that developing progenitors and precursor cells perform macropinocytosis, but also that macropinocytosis is essential for their appropriate lineage commitment. Indeed, our data suggest a model whereby growth factor-induced macropinocytosis determines the ultimate outcome of the differentiation event, in contrast to a model where independent competing growth factors dictate the outcome of progenitor differentiation. Our proposed model raises an important question: if M-CSF can drive progenitor or precursor cell differentiation into either lineage depending on whether macropinocytosis is induced or not, what are the distinct signals or factors that arise during macropinocytosis that drive one fate over another? The answer may relate to signaling in or at the macropinosome itself. For instance, amplification of growth factor signaling strength by macropinosome cup formation (Huynh et al., 2012; Kay et al., 2018) may inform cell fate determination. Alternatively, solutes in the macropinosome itself could relay requisite signals to differentiate into one lineage over another. Solute-sensing signaling hubs, such as the mechanistic target of rapamycin (mTOR) have been previously shown to respond to growth factor presence in the macropinosome (Yoshida et al., 2015). Finally, although we demonstrate that WNK1 is required for M-CSF-induced macropinocytosis, it remains unknown if WNK1 is directly responsible for lineage commitment (mediator) or is instead indirectly required via its role in regulating macropinocytosis (moderator). Regardless, determining the specific signal(s) arising in or at the macropinosome that dictate progenitor and precursor cell lineage commitment will be an important area of work moving forward.

## STAR Methods

### Contact for Reagent and Resource Sharing

Requests for resources and reagents should be directed to the Lead Contact, Justin S. A. Perry.

### Mice

Animals were housed at the Memorial Sloan Kettering Cancer Center (MSKCC) animal facility under specific pathogen free (SPF) conditions on a 12-hour light/dark cycle under ambient conditions with free access to food and water. WNK1 deletion from myeloid cells was achieved by crossing *Csf1r^iCre^* (Jackson Laboratories, Strain #021024) mice with Wnk1^fl/fl^ mice (a gift from Chou-Long Huang, University of Iowa, as previously described (Mayes-Hopfinger et al., 2021)), to obtain wildtype (WT; *Csf1r^Cre–^; Wnk1^fl/fl^*), heterozygous (*Csf1r^Cre+^; Wnk1^fl/+^*), and conditional WNK1 KO littermates (*Csf1r^Cre+^; Wnk1^fl/fl^*). For *Csf1r^Cre+^; Wnk1^fl/fl^* studies, animals were weighed and monitored to determine failure to thrive. For adoptive monocyte transfer experiments, CD45.1+ congenic mice were obtained from Jackson Laboratories (Strain #0002014). All studies were approved by the Sloan Kettering Institute (SKI) Institutional Animal Care and Use Committee (IACUC).

### Tissue collection and processing

Mice between 3 to 4 weeks of age were euthanized and perfused using a 23G needle with 20 mL of cold PBS without Ca^2+^ or Mg^2+^ (Corning, 21-040-CM) with 5 mM EDTA (Invitrogen, 15575-038). Blood was collected immediately following euthanasia, prior to perfusion via cardiac puncture, and transferred to heparin-coated tubes (Grenier, 450474). Brain, lungs, liver, kidneys, spleen, and heart were dissected, weighed, and minced into small fragments, and incubated at 37°C for 30 min in enzyme mix consisting of PBS with 160 Wünsch U/ml collagenase D (Sigma-Aldrich, 11088866001), 5% heat-inactivated FBS (Sigma, 1306C), and 10 mM HEPES (Gibco, 15630-080) (‘PFH’ buffer). The digestion reaction was stopped by incubation with 10 mM EDTA for 5 min. After enzyme digestion, all tissues were further dissociated by mechanical disruption using 70 µm cell strainers and an 18G needle with 3-mL syringe plunger in 6-well plates containing 3 mL of cold PFH buffer. Single-cell suspensions were transferred to 15 ml tubes and pelleted by centrifugation at 1500 rpm for 5 min at 4°C. Cell pellets were then resuspended in 38% Isotonic Percoll (10% 10X PBS with 90% Percoll (GE Healthcare, 17-0891-01)) in PFH buffer to gradient separate leukocytes by spinning at 2000 rpm for 30 min at RT with no brake. Bone marrow cells were collected by flushing femurs and tibias using a syringe and a 25G needle with 10 mL PFH buffer. Bone marrow was dissociated by gently flushing through a 70 μM filter with the back of the syringe plunger. Collected blood, spleen (post-digestion), and bone marrow suspensions were spun down and lysed with 1 mL red blood cell lysis buffer (Sigma, R7757) for 5 min, washed with 10 mL PFH buffer and centrifuged at 1500 rpm for 5 min at 4°C.

### Flow cytometry

Isolated leukocytes were washed once in PBS and stained with Live/Dead Fixable Aqua stain (Thermo Fisher Scientific, L34966) diluted 1:1000 in PBS (no FBS) for 30 min at RT, protected from light. Samples were then washed once in PFH buffer and resuspended in 50 μl of PFH containing purified anti-mouse CD16/32 antibody (1:50, BioXCell, BE0307) and incubated for 10 min on ice. Samples were then directly incubated with fluorochrome-conjugated antibodies for an additional 15 min on ice, then washed in PFH buffer and analyzed by flow cytometry using an Attune NxT flow cytometer (Invitrogen). In all tissues, live single cells were gated by exclusion of Live/Dead Aqua-positive dead cells, positive gating of side scatter and forward scatter, and doublet cell exclusion using forward scatter width against forward scatter area, as previously described (Gomez Perdiguero et al., 2015). To calculate cell numbers per sample, counting beads (Thermo Fisher Scientific, C36950) were added at known concentrations to known volumes of samples and gated for back calculation after flow cytometry analysis. All post-acquisition analysis was conducted using FlowJo 10.7.1 (BD Biosciences).

### Microscopy

#### Tissue Processing

Tissues were harvested from mice and immediately submerged in 4% formalin (Stat-Labs, 28530-1) for 24-48h at 4°C. Tissues were subsequently washed 3x in PBS without Ca^2+^ or Mg^2+^ (Corning, 21-040-CM), then submerged in 30% sucrose (Sigma, S0389) in PBS overnight or until they sunk. Tissues were then embedded in Tissue-Tek OCT compound (Sakura, 4583), snap-frozen on dry ice, and immediately stored at −80°C until sectioning. 10 μm sections were cut on a Leica Microm HM550 and mounted on charged slides (Fisherbrand Superfrost Plus Gold, 15-188-48) for H&E and IF staining.

#### H&E staining

Briefly, tissue sections were brought to RT and dehydrated in 95% ethanol, stained with Harris Modified Hematoxylin (Fisher Scientific, SH26-500D) for 5 min, run under tap water until optimal bluing was achieved, then submerged into 0.5% acid alcohol for 10 s to stop the reaction. Sections were subsequently dehydrated in 95% ethanol and submerged in Eosin Y (Sigma-Aldrich, 1732-87-1) for 30 s. Sections were then treated with increasing concentrations of ethanol (70%, 95%, 100%), cleared with xylenes (Sigma-Aldrich, 534056), coverslip-mounted with Permount (Fisher-Scientific, SP15-100), and allowed to dry for 24h prior to Differential Interference Contrast (DIC) imaging.

#### DIC Imaging and Analysis

H&E and brightfield images were taken on an EVOS M5000 imaging system (Invitrogen) at 10X, 20X, and 40X magnification. Images were blinded, processed, and analyzed using Fiji (‘Fiji is Just Image J’, NIH) by first converting images to binary and applying the “analyze particle” plugin for vesicle counting and characterization. Cell outlines were drawn free-hand and size, area, and circularity were measured on 40X brightfield images for all mouse and human cell culture images.

#### Immunofluorescence

Tissue sections were brought to RT and rehydrated in PBS for 5 min, then blocked and permeabilized in 10% normal goat serum (Sigma-Aldrich, G9023) and 0.1% Triton-X-100 (Sigma-Aldrich, T9284) for 2h. Individual sections were then separated using a barrier pen (Vector Labs, H-4000) and incubated in primary antibodies at 4°C overnight in a humidified chamber as follows: neutrophil staining, anti-rat Ly6G-PE, 1:200 (1A8, BioLegend, 127607), macrophage staining, anti-rabbit Iba1, 1:500 (Polyclonal, Wako/Fujifilm, 019-19741). Slides were then washed 3x for 5 min in PBS. Sections were subsequently stained with IgG H+L secondary antibodies at room temperature for 1h, using Goat-anti-rat-AF568 (Ly6G) and Goat-anti-rabbit-AF647 (Iba1) (Life Technologies/Invitrogen, A11077 or A23733) diluted 1:500. Finally, sections were washed 3x in PBS with the second to last wash containing nuclei dye Hoechst (Thermo Fisher, H3570) diluted 1:1000. Slides were mounted with coverslips and Prolong Gold Antifade (Thermo-Fisher P36934), sealed with clear nail polish, and allowed to dry overnight before imaging. Images (40X) were taken as Z-stacks on an inverted Zeiss LSM 980 with Airyscan 2 and processed/analyzed in Fiji.

### Isolation and culture of mouse bone marrow monocytes (precursor cells)

Monocytes from 4-week-old Cre− and Cre+ mice were obtained by flushing femurs and tibias with 10 mL DMEM (Fisher Scientific, MT10017CV) containing 20% FBS. To enrich for Ly6C+ monocytes, total bone marrow was subjected to the mouse monocyte isolation kit (MACS Miltenyi Biotec, 130-100-629), performed according to manufacturer instructions. Resultant monocytes were collected, counted, and cultured in αMEM (Corning, 15-012-CV) supplemented with 10% FBS and 1% PSQ, with 50 ng/mL recombinant murine M-CSF (PeproTech, 315-02) at 37 °C and 5% CO_2_ in a 12-well plate with vehicle or WNK463 (10 μM). Cells were monitored daily via fluorescent microscopy (EVOS 5500) and collected after 7d of differentiation. Differentiated monocytes were then either harvested for immediate flow cytometry or exposed to indicated treatments prior to analysis.

### Isolation and culture of mouse bone marrow progenitors

Progenitors from 4-week-old Cre− and Cre+ mice were obtained by flushing femurs and tibias with 10 mL DMEM (Fisher Scientific, MT10017CV) containing 20% FBS. To enrich for lineage-negative cells, total bone marrow was subjected to the mouse lineage depletion kit (MACS Miltenyi Biotec, 130-090-858), performed according to manufacturer instructions. For imaging studies, prior to progenitor isolation, 24 well TC-treated plates or 8 well glass chambers (Lab-Tek II, 155409) were coated for 1h at RT with Poly-L-lysine hydrobromide (Sigma, P2636) at 0.5mg/mL in water, washed 3x with water, and subsequently coated overnight at 4°C with 2.5μg/mL recombinant human Laminin (Thermo Fisher Scientific, A29249) in PBS. The following day, resultant progenitor lineage-negative cells were collected, counted, and cultured in DMEM containing 20% heat-inactivated FBS, 50 ng/mL murine stem cell factor (PeproTech, 250-03), 10 ng/mL recombinant murine IL-3 (PeproTech, 213-13), and 20 ng/mL recombinant human IL-6 (PeproTech, 200-06) without PSQ for 48-72h at 37 °C and 5% CO_2_ (modified from (Wilkinson et al., 2020)). Prior to stimulation with M-CSF or various inhibitors (see macropinocytosis assays and intracellular chloride analysis), cells were serum starved for 1h. Images were captured 1h after initiation unless otherwise stated. Differentiated progenitors were then either harvested for immediate flow cytometry and western blot analysis or exposed to indicated treatments prior to analysis.

### Adoptive transfer experiments

Bone marrow monocytes or progenitors were isolated from 8-to-12-week-old CD45.1 congenic male and female donors. To enrich for Ly6C+ cells (monocytes), total bone marrow was stained with Ly6C-PE (BD Pharmingen, 560592) for 10 min on ice, washed 1x with PBS, then incubated with anti-PE microbeads from the anti-PE Isolation Kit for mouse (MACs Miltenyi Biotec, 130-048-801), used as suggested by the manufacturer. For progenitors, total bone marrow was incubated with a cocktail of biotinylated antibodies against a panel of lineage antigens (CD5, CD45R (B220), CD11b, Anti-Gr-1 (Ly6C/G), 7-4, and Ter-119 antibodies) and subsequently directly incubated with anti-biotin microbeads from the lineage cell depletion kit for mouse (MACs Miltenyi Biotec, 130-090-858), used as suggested by the manufacturer. Total Ly6C+ or Lin-cells were counted using Cell Counter chamber slides (Invitrogen, C10283) in a Countess II FL (Invitrogen). Approximately 1×10^6^ CD45.1+ Ly6C+ or 1×10^5^ lin-cells were diluted in sterile PBS prior to intraperitoneal injection. Injections occurred at P5, P8, and P11 or P2, P5, P8 as indicated. Mice were either monitored for extended survival or sacrificed at P28 (2-3 weeks after final injection). Denoted organs/tissues were analyzed for the fraction of CD45.1+ cells (chimerism) in the total CD45+ pool. Alternatively, CD45.1 congenic mice were used as wildtype recipients for bone marrow transplantation experiments where cage mates were treated with 1 dose of 900rad lethal irradiation and put on broad-spectrum antibiotics diluted in drinking water at 2mg/mL for 1 week following irradiation (Baytril 100, Enrofloxacin, Bayer). Irradiated animals were IP injected with 5×10^5^ lin-cells obtained from Cre− or Cre+ mice as described above within 24h of lethal irradiation.

### Isolation and culture of human blood monocytes (precursor cells)

Peripheral blood mononuclear cells (PBMCs) isolated from deidentified healthy blood donors were obtained by gradient separation using 1-Step Polymorphs (Accurate Chemical, AN221725). CD14+ monocytes were isolated with Mojosort human CD14 nanobeads (BioLegend, 480093). 5×10^6^ monocytes were seeded and differentiated into macrophages for 7d in RPMI-1640 (Corning, 17-105-CV) supplemented with 10% FBS and 1% PSQ, with 50 ng/mL recombinant human M-CSF (PeproTech, 300-25) at 37 °C and 5% CO_2_. Human monocyte-derived macrophages (HMDMs) were re-seeded at 2×10^5^ cells per mL (1 mL per well) in a 12-well plate with vehicle or WNK463 (10 μM). Cells were monitored daily via fluorescent microscopy (EVOS 5500) with images presented in the manuscript collected after 7d of differentiation. HMDMs were then harvested by scraping cells directly into media, centrifuged at 2500 rpm in a benchtop centrifuge for 5 min at RT, incubated with FC block and indicated antibodies, then analyzed by flow cytometry (see **Table C** for human antibodies and dilutions).

### Human pluripotent stem cell (hPSC) cultures

Deidentified human PBMCs were reprogrammed into pluripotent stem cells (PSCs) using recombinant Sendai viral vectors that express Yamanaka factors (Yang et al., 2008). PSCs were maintained on CF1 mouse embryonic fibroblasts (ThermoFisher, A34181) in ESC medium (ThermoFisher, 10829018) supplemented with bFGF (PeproTech, 100-18B). Embryoid bodies (EBs) were formed by disassociating and maintaining PSCs in ESC media supplemented with a Rho-associated protein kinase (ROCK) inhibitor (10μM; Sigma, Y0503) while orbitally shaking (100 rpm) for 6d (Lachmann et al., 2015). Mature EBs were transferred to tissue culture-treated 6-well plates and maintained in APEL media (Stem Cell Technologies, 5270) supplemented with 5% PFHM-II (Fisher, 12040077), 1% penicillin-streptomycin, human IL-3 (25ng/mL; PeproTech, 200-03) and M-CSF (50 ng/mL; PeproTech, 300-25), in the presence of either vehicle or WNK463 (10 μM) for an additional 7d. In some experiments, myeloid progenitors were derived normally, then treated with M-CSF in the presence of vehicle or WNK463 (10 μM) for an additional 5-10d to produce terminally-differentiated macrophages. Resultant PSC-derived macrophages were then analyzed by fluorescent microscopy or flow cytometry similar to analyses performed with HMDMs.

### Western Blotting

Mature macrophages or progenitors were seeded in 12-well plates at a density of 2.5×10^5^ cells per well unless otherwise stated. In some experiments, technical replicate wells were pooled to increase protein concentration per condition. After treatment with cytokines and/or inhibitors, cells were washed 1x in HBSS without Ca^2+^ or Mg^2+^(Gibco, 14170-112) as phosphate-containing buffers can disrupt phosphorylation assay data. Pelleted cells were lysed in Pierce RIPA buffer (Thermo scientific, 89901) containing protease and phosphatase inhibitor cocktails (Roche, 04 906 845 001). Lysates were then used in total-protein western blots (Nu-Page 4-12% Bis-Tris Gel, Invitrogen, NP0335BOX)) or phosphorylation blots (6 or 7.5% Phos-Tag gels, Wako/Fujifilm, 192-18001). Protein blots were transferred to PVDF membranes (BioRad, 1620177) using the Trans-Blot Turbo System (BioRad, 170-4270) according to the manufacturer protocol. Membranes were blocked for 30 min with 3% BSA in TBS-T (TBS: Teknova, T1680, Tween-20: Sigma, P1379), washed 3x, then probed overnight at 4 °C with anti-WNK1 (R&D Systems, AF2849) and anti-β-actin-HRP (Santa Cruz, sc47778 HRP) or anti-transferrin receptor (TFR; Abcam, ab2140398) antibodies. Antibodies were diluted in TBS-T containing 3% BSA. Secondary goat, rabbit, and mouse antibodies directly linked to HRP (Amersham) were diluted at 1:5000 in 3% BSA in TBS-T and added to membranes for 1h at RT. Membranes were washed 3x and treated with SuperSignal West Pico Plus ECL (Thermo scientific, 34578) for 5 min in the dark. Membranes were imaged using the iBright 1500 imaging system (Invitrogen) to analyze chemiluminescence. Densitometry was calculated using Fiji and expression was normalized as a ratio of the individual phosphorylated bands to the total protein band, all divided by the total protein of the loading control (unless otherwise stated).

### Macropinocytosis assays

To assess macropinocytosis, the reagents DQ-Red BSA (Thermo Fisher Scientific, D12051) or Tetramethylrhodamine (TMR)-dextran, 70,000 MW, lysine fixable (Thermo Fisher Scientific, D1818) was added to indicated cells at a concentration of 10 μg/mL of DQ-Red BSA or 1 mg/mL TMR-dextran for 1h. Cells were then washed 1x with PBS and analyzed via fluorescent microscopy (EVOS M5000 or Zeiss LSM 980) or flow cytometry. In some experiments, indicated cells were collected and lysed for western blot. To inhibit macropinocytosis, the compounds 5-[ethyl(1-methylethyl) amino]-2-pyrazinecarboxamide (EIPA, at 10 μM, where indicated, Tocris, 1154-25-2), LY294002 (50 μM, SelleckChem, S1105), or Apilimod (10 μM, MedChemExpress, HY-14644) were added 1h prior to macropinocytosis induction.

### Analysis of intracellular chloride

Prior to the initiation of macropinocytosis assays, lineage-negative progenitors were labeled with *N*-(Ethoxycarbonylmethyl)-6-Methoxyquinolinium Bromide (MQAE; Molecular Probes, Thermo Fisher Scientific, E3101) for 1h in stem cell-specific DMEM without serum. Cells were then washed 1x and treated with M-CSF (50 ng/mL) for an additional hour. In some experiments, the chloride ionophore tributyltin chloride (TBTC; Sigma-Aldrich, 45713-250MG) was added (10 μM) in the presence of vehicle or WNK463 (10 μM). MQAE, when excited, emits optimally at 460 nm and can be detected via flow cytometry analysis. Furthermore, MQAE is insensitive to both pH and bicarbonate, making it optimal for detecting chloride changes irrespective of other physiological changes occurring in the cell (Perry et al., 2019).

### Statistics and reproducibility

Statistical analyses were performed using GraphPad Prism 9.3.1. Significance was determined using Kaplan-Meier’s Log-Rank Mantel-Cox test, independent samples Student’s *t*-tests, or one-way ANOVA, according to data requirements. All experiments were performed independently at least three times except for those detailed below. The experiments and analyses outlined in **Figure 4** were performed independently twice but consisted of individual analysis of at least 100 unique cells for each independent replicate. Furthermore, these analyses were performed on a subset of experiments that were performed more than three times. The macropinocytosis assays performed in **Figure 5E** were performed across two independent experiments but included 2 independent Cre− mice and 2 independent Cre+ mice.

## Code Availability

No unique code was generated or used in the preparation of this manuscript.

## Acknowledgements

We thank members of the Perry laboratory and co-authors for edits and discussions related to this manuscript. We also thank Lydia Finley (MSKCC) for critical analysis of the data and experiments, the New York Blood Center for human total blood, and the MSKCC Stem Cell Research Core for human pluripotent stem cell derivation. This work was supported by grants to J.S.A.P. from the NIH (NCI 5R00CA237728; NIGMS 1DP2GM146337), a Pew Biomedical Scholars Award, a pilot award from the CSBC Research Center for Cancer Systems Immunology at MSKCC (NCI 5U54CA209975), training grant support to A.J.T. from the NIH (NCI 5T32CA009149), to C.-L.H. from the NIH (NIDDK DK111542) and MSKCC Cancer Center Support Grant P30CA008748. Some figure panels were created using the commercial version of BioRender.

## Contributions

A.J.T. and J.S.A.P. planned, with A.J.T. executing, the majority of the experiments. W.S.R., Z.W., P.S., Z.L., I.C.M., and Y.-T.W. assisted with the performance and analysis of *in vivo*/ex vivo experiments. G.Z. assisted with *in vitro* microscopy experiments. J.X. and C.-L.H. generated *Wnk1*^fl/fl^ mice. M.O. assisted with the planning and execution of macropinocytosis assays. A.S.K. and M.S.G. assisted with human PSC experiments. C.N.P., T.V., and C.D.L. assisted with the analysis and interpretation of *in vivo*/ex vivo experiments. All authors assisted in the preparation and review of the manuscript.

