## Supplemental Figures for "WNK1 enforces macrophage lineage fidelity"

**Figure S1. *Csf1r*$^{Cre+}$*;Wnk1*$^{fl/fl}$ mice do not follow Mendelian inheritance.**

**(A)** Prevalence of homozygous wildtype (white), heterozygous (fl/+, orange), and homozygous WNK1-deficient mice (fl/fl, blue) per litter.

**(B)** Percentage of genotype in each litter. Shown are homozygous wildtype (white), heterozygous (fl/+, orange), and homozygous WNK1-deficient mice (fl/fl, blue). Data shown as mean ± SEM. Statistical significance was determined via one-way ANOVA. ns = not significant. ****$p < .0001$.

**Figure S1: *Csf1r*$^{Cre/+}$*;Wnk1*$^{fl/fl}$ mice do not follow Mendelian inheritance**

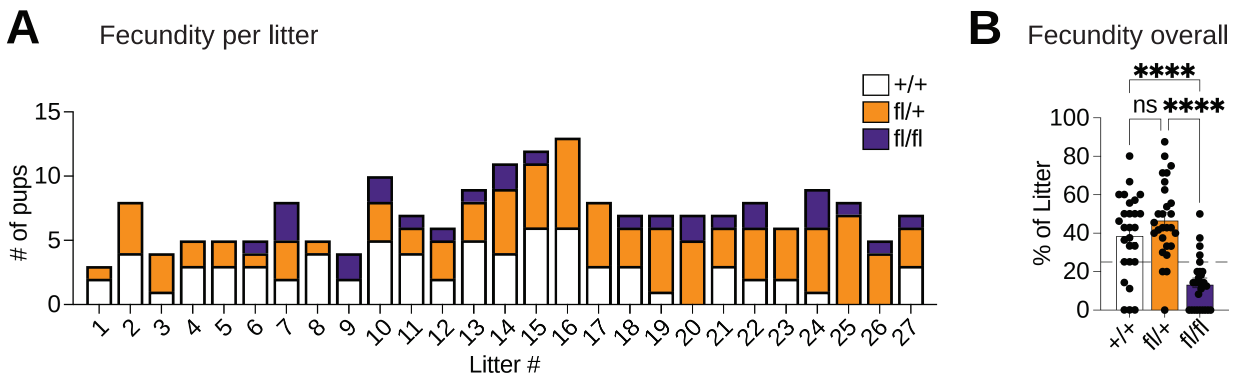

**Figure S2. Tissue-resident macrophages are absent in many organs of *Csf1r*$^{Cre+}$*;Wnk1*$^{fl/fl}$ mice.**

**(A)** Flow cytometry gating scheme for **Figures 1** and **2** and **Figures S2** and **S7** using lung as an example.

**(B-E)** Data are from experiments performed in **Figure 1**. Shown are H&E, flow cytometry, and immunofluorescence analyses of additional essential tissues obtained, analyzed, and presented as in **Figure 1C-F**. Data are from at least four *Csf1r*$^{Cre+}$;*Wnk1*$^{fl/fl}$ mice and five littermate controls. Statistical significance was determined via independent samples *t*-test. ns = not significant. *$p < .05$. **$p < .01$. ***$p < .001$. ns = not significant.

**(F)** Flow cytometry analysis of CX3CR1+ monocytes isolated from experiments performed in **Figure 1**. Shown are representative flow cytometry plots (top) and summary plots of absolute numbers per milligram of tissue and frequencies (bottom) of $CX_3CR1$+ monocytes. Data are from six *Csf1r*$^{Cre+}$;*Wnk1*$^{fl/fl}$ mice and six littermate controls. Data shown as mean ± SEM. Statistical significance was determined via independent samples *t*-test. *$p < .05$. **$p < .01$, ns = not significant.

#### Figure S2: Tissue resident macrophages are absent in essential organs

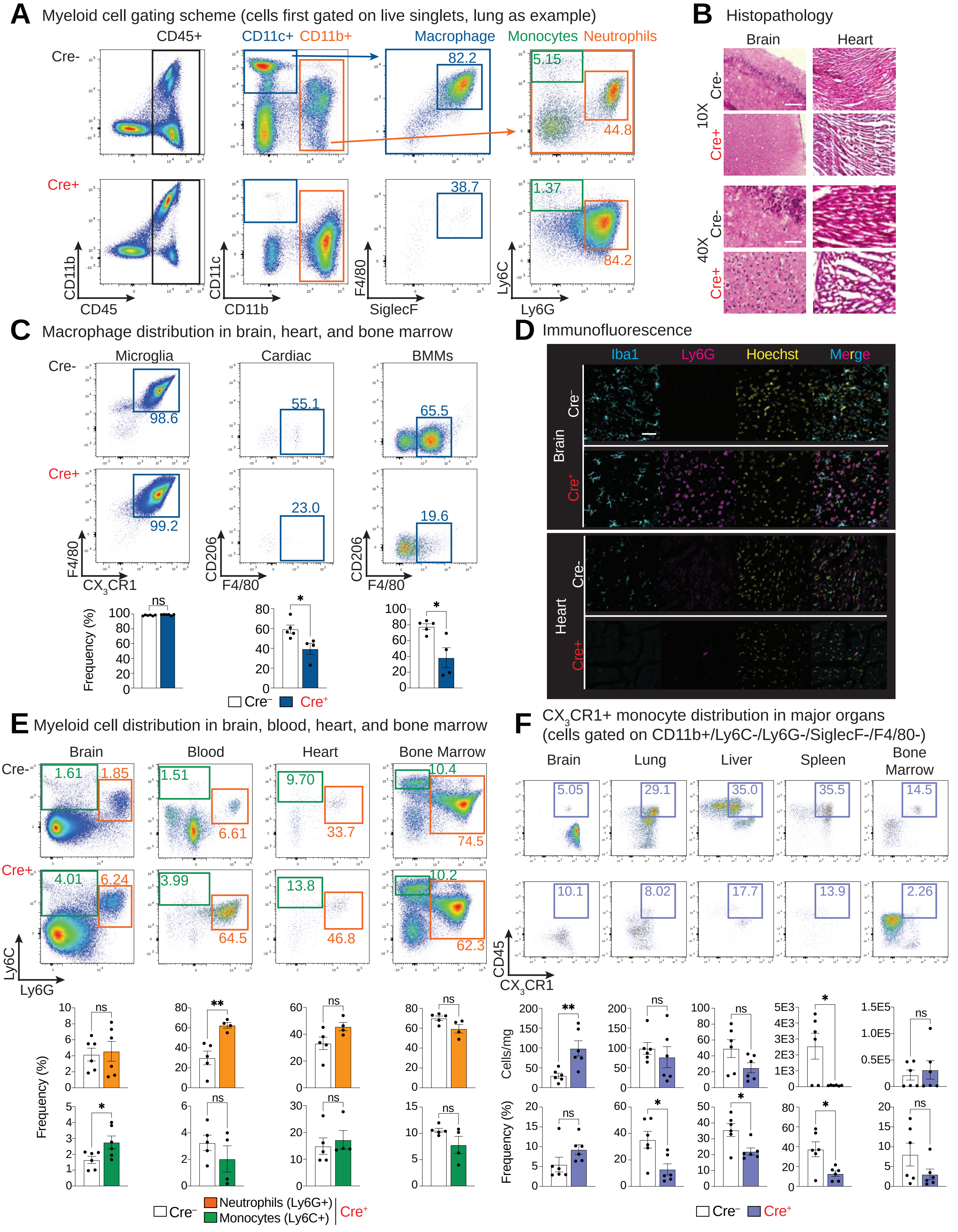

**Figure S3. Lymphoid cell distribution in major organs of *Csf1r*$^{Cre+}$*;Wnk1*$^{fl/fl}$ mice.**

**(A-C)** Flow cytometry analysis of the lymphoid compartment isolated from experiments performed in **Figure 1** and **S2**. Shown are summary plots of absolute numbers per milligram of tissue and frequencies of (**A**) B cells (CD19+ MHCII+ CD4- CD8- CD11b- CD45+), (**B**) CD4+ T cells (CD4+ CD8- CD19- CD11b- CD11c- CD45+), and (**C**) CD8+ T cells (CD8+ CD4- CD19- CD11b- CD11c- CD45+). Data are shown as mean ± SEM. Data are from six *Csf1r*$^{Cre+}$;*Wnk1*$^{fl/fl}$ mice and six littermate controls. Statistical significance was determined via independent samples *t*-test. *$p$ < .05, **$p$ < .01, ns = not significant.

**Figure S3: Lymphoid cell distribution in major organs**

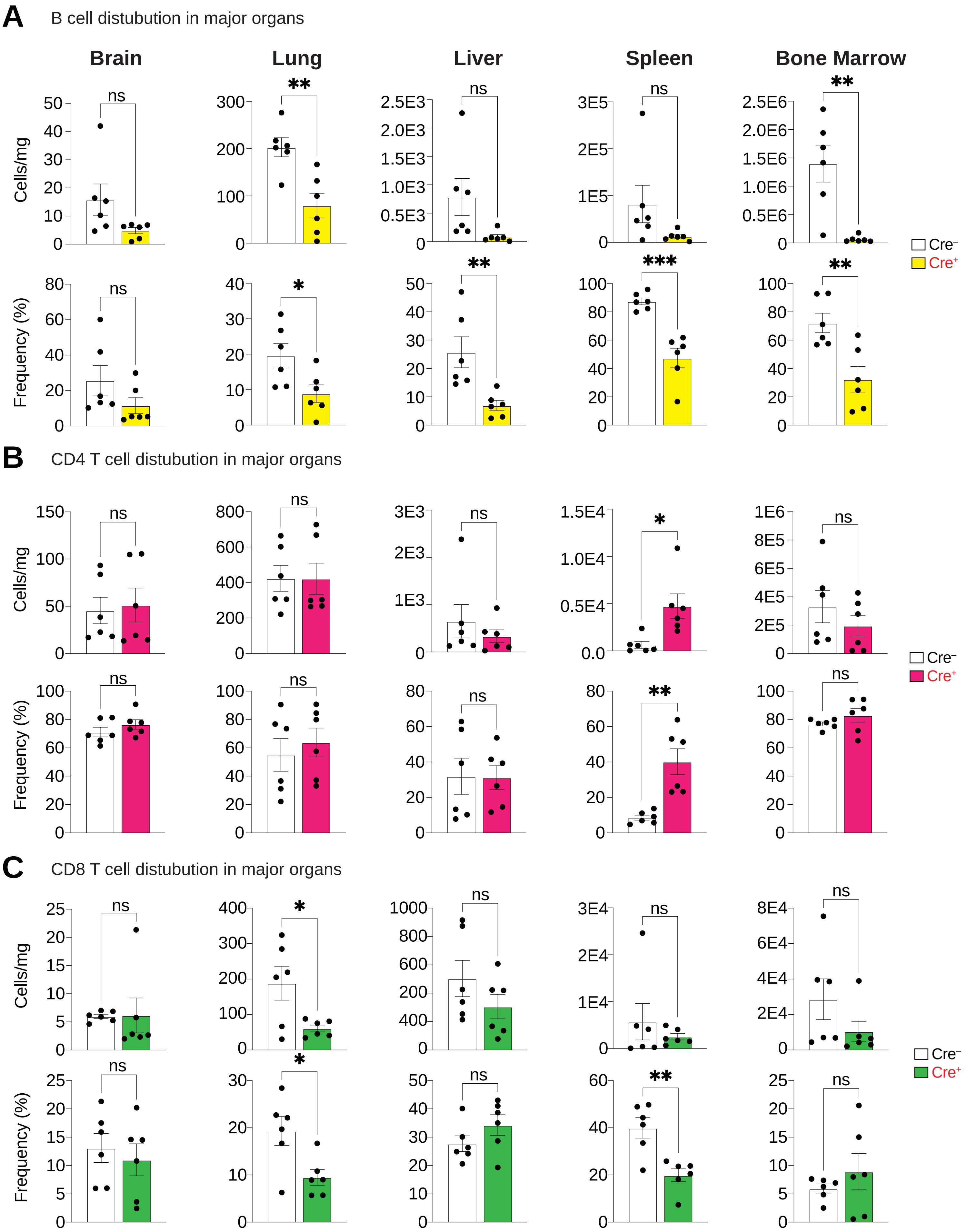

**Figure S4. *Csf1r*$^{Cre+}$*;Wnk1*$^{fl/fl}$ mice exhibit skewed myelopoiesis.**

**(A)** Schematic of hematopoiesis of long-term (LT), short-term (ST), and multipotent progenitors (MPPs) with corresponding markers. Also shown is the figure panel that corresponds to the lineage state analyzed. Where possible, bar graph color corresponds to cell color in the schematic.

**(B)** Quantification of hematopoietic stem cell (HSC) cKit and Sca-1 expressing populations. Representative flow cytometry (left) and summary (frequency and cell counts; right) plots of cKit and Sca-1 expressing HSC populations. Bone marrow cells were first gated on lineage negative (lin-) cells, then analyzed for cKit and Sca-1 positivity. Data are from n=7 *Csf1r*$^{Cre+}$*;Wnk1*$^{fl/fl}$ (Cre+) or n=7 *Csf1*$^{Cre-}$*;Wnk1*$^{fl/fl}$ (Cre-) mice across four independent experiments. Data are shown as mean ± SEM. Statistical significance was determined via independent samples *t*-test. $^{**}p < .01$, ns = not significant.

**(C)** Quantification of LT-HSC, ST-HSC, and early MPP populations. Representative flow cytometry (left) and summary (frequency and cell counts; right) plots of the HSC subtypes long-term (LT: CD48- CD150+, blue), short-term (ST: CD48- CD150-, orange), and multipotent progenitor (MPP: CD48+ CD150-, green). Data are from n=7 Cre- and n=7 Cre+ mice across four independent experiments. Data are shown as mean ± SEM. Statistical significance was determined via independent samples *t*-test. * $p < .05$, $^{**}p < .01$, $^{***}p < .001$, ns = not significant.

**(D)** Quantification of early and late stage MPPs. Representative flow cytometry (left) and summary (frequency and cell counts; right) plots of Flt3 negative (Flt3$^{-}$), early (Flt3$^{low}$), and late (Flt3$^{high}$) MPPs. Cells were initially gated as CD48+ CD150-. Data are from n=7 Cre- and n=7 Cre+ mice across four independent experiments. Data are shown as mean ± SEM. Statistical significance was determined via one-way ANOVA with Tukey post hoc test. * $p < .05$, $^{**}p < .01$, $^{***}p < .001$, ns = not significant.

**(E)** Quantification of MPP subsets. Representative flow cytometry (left) and summary (frequency and cell counts; right) plots of the following MPP subsets: MPP2 (erythrocyte/megakaryocyte lineage, orange bar), MMP3 (granulocyte/monocyte/macrophage lineage, blue bar), and MPP4 (lymphoid lineage, purple bar). Data are from n=7 Cre- and n=7 Cre+ mice across four independent experiments. Data are shown as mean ± SEM. Statistical significance was determined via independent samples *t*-test. * $p < .05$, $^{***}p < .001$, ns = not significant.

#### Figure S4: Cell distribution during stages of hematopoiesis

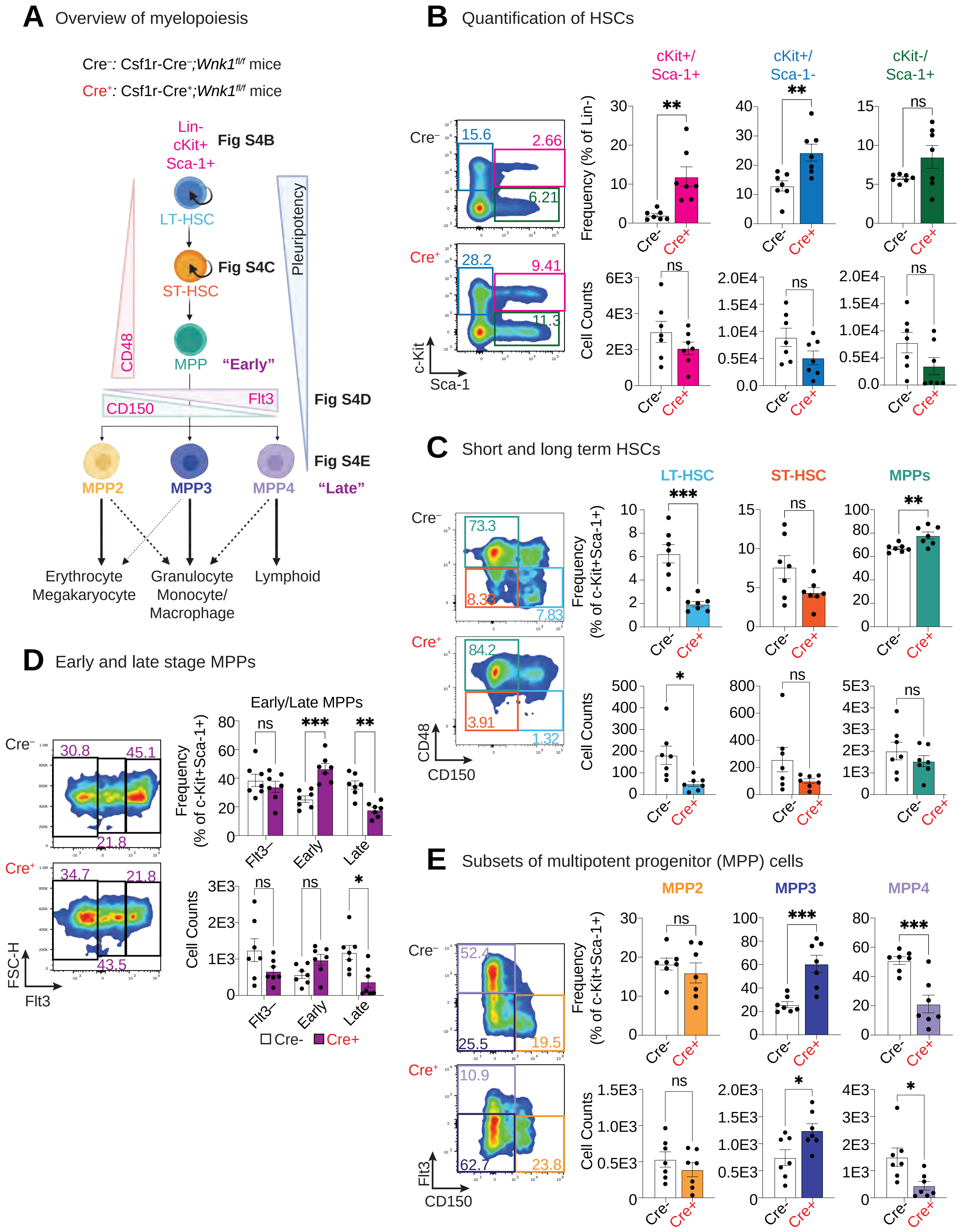

**Figure S5. CD45.1 donor cell contribution in Cre+ recipient mice.**

**(A)** Representative flow cytometry plot of the brain (left) and lungs (right) from Cre+ recipient mice (CD45.2+) injected with CD45.1+ monocytes as in **Figure 2B**.

**(B-D)** Analysis of **(B)** neutrophil, **(C)** Ly6C+ monocyte, and **(D)** $CX_3CR1$+ monocyte tissue frequencies from monocyte transfer experiments (from **Figure 2B**) or progenitor transfer experiments (from **Figure 2C**). Shown are summary plots of cell frequency as a fraction of CD45.1+ CD11b+ cells in the brain, lungs, kidney, liver, and spleen of Cre- (white), Cre+ (light shaded), and Cre+ mice injected with wildtype CD45.1+ monocytes or progenitors (dark shaded). Data are from two independent experiments with three (Cre-), two (Cre+ progenitors), or three (Cre+ monocytes) mice per group. Data are shown as mean ± SEM. Statistical significance was determined via one-way ANOVA. *$p < .05$, **$p < .01$, ***$p < .001$, ****$p < .0001$, ns = not significant.

**Figure S5: CD45.1 donor cell contribution in Cre⁺ recipient mice**

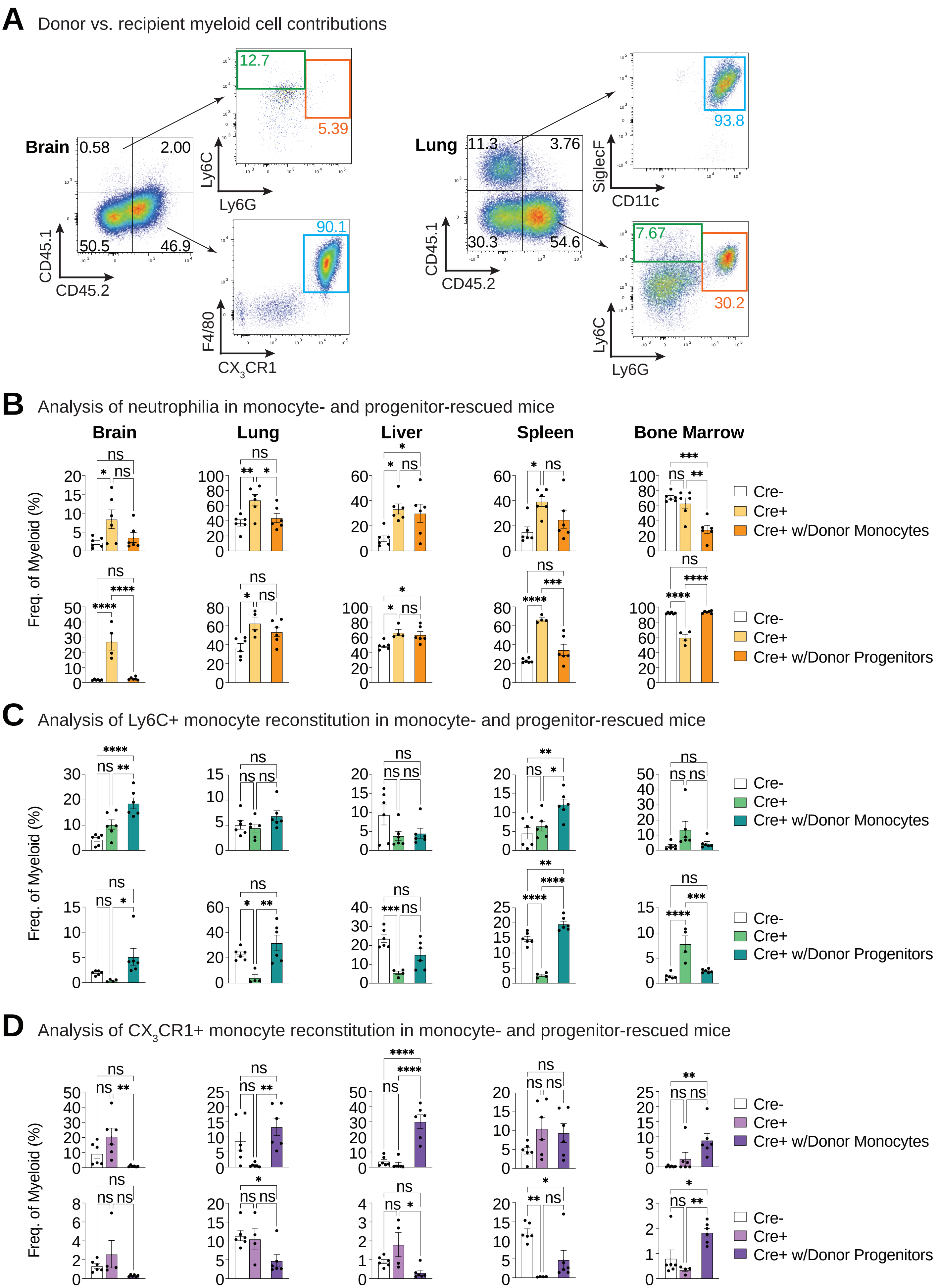

**Figure S6. *Csf1r*$^{Cre+}$*;Wnk1*$^{fl/fl}$ cells do not become canonical macrophages.**

**(A)** Shown is the experimental strategy used to test macrophage differentiation of bone marrow myeloid progenitors or monocytes from *Csf1r*$^{Cre+}$*;Wnk1*$^{fl/fl}$ (Cre+) or *Csf1*$^{Cre-}$*;Wnk1*$^{fl/fl}$ (Cre-) mice in response to M-CSF.

**(B, C)** Representative flow cytometry plots and summary data of CSF1R, CD206, CD11b, CD11c, F4/80, Ly6C, and MHCII in monocytes (**B**) or myeloid progenitors (**C**) treated with M-CSF as in (**A**). Data are from three independent experiments and shown as mean ± SEM. Statistical significance was determined via independent samples *t*-test. *$p < .05$, **$p < .01$, ***$p < .001$, ns = not significant.

**Figure S6: *Csf1r$^{Cre/+}$:Wnk1$^{fl/fl}$* progenitors do not become canonical macrophages**

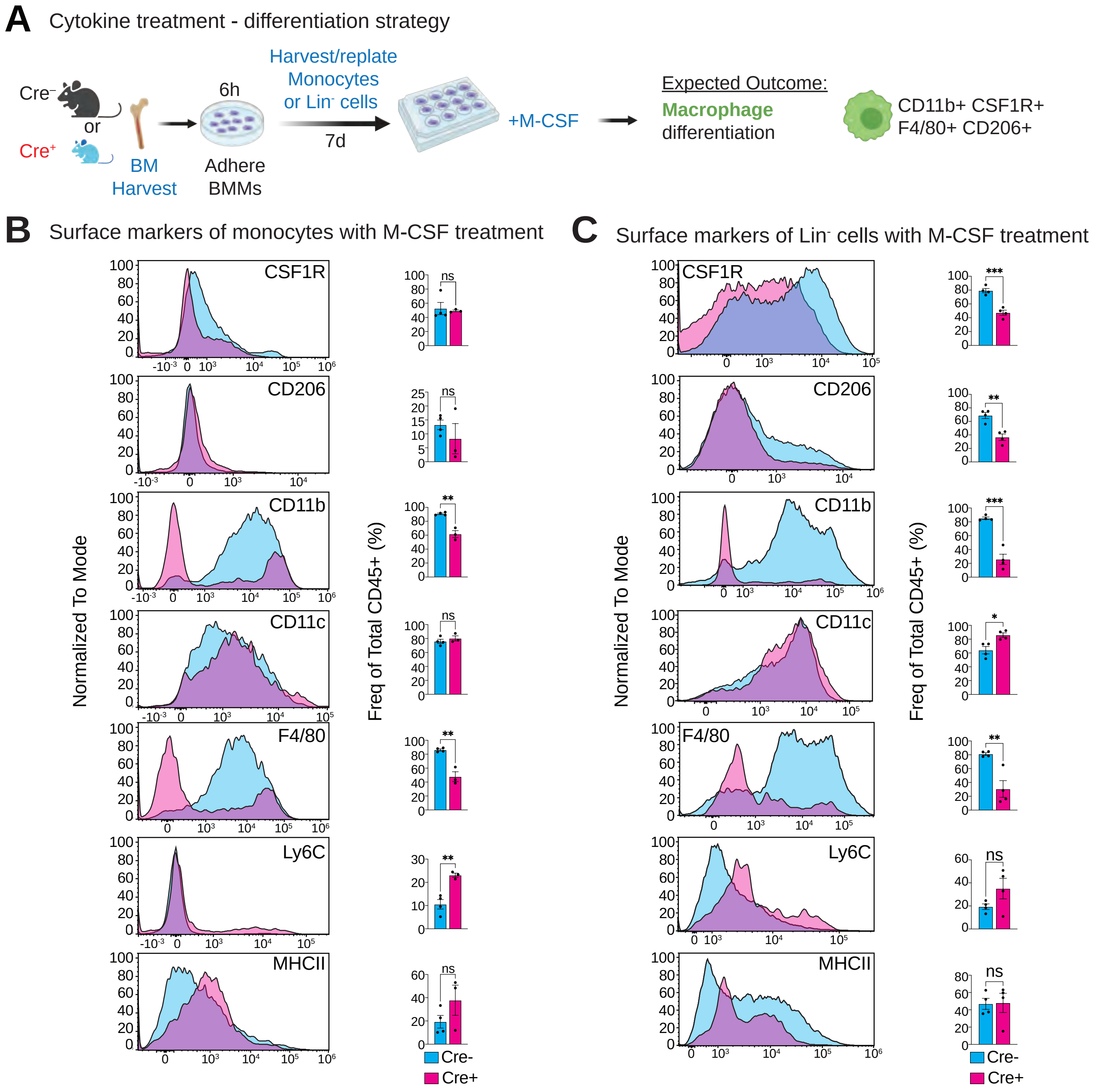

**Figure S7. Myeloid cell frequencies in bone marrow transplantation experiments.**

**(A-D)** Analysis of **(A)** macrophages, **(B)** neutrophils, **(C)** Ly6C+ monocytes, and **(D)** $CX_3CR1$+ monocytes arising from Cre- (Cre- donor progenitors into wildtype adult mice; white) and Cre+ (Cre+ donor progenitors into wildtype adult mice; shaded) across tissues analyzed. Data are from two independent experiments, with three (Cre+, n=6 total) or two (Cre-, n=4 total) mice per group. Data are shown as mean ± SEM. Statistical significance was determined via independent samples *t*-test. *$p$ < .05, **$p$ < .01, ns = not significant.

### Figure S7: Bone marrow transplantation myeloid cell frequencies

**A** Analysis of TRM frequencies in irradiated mice

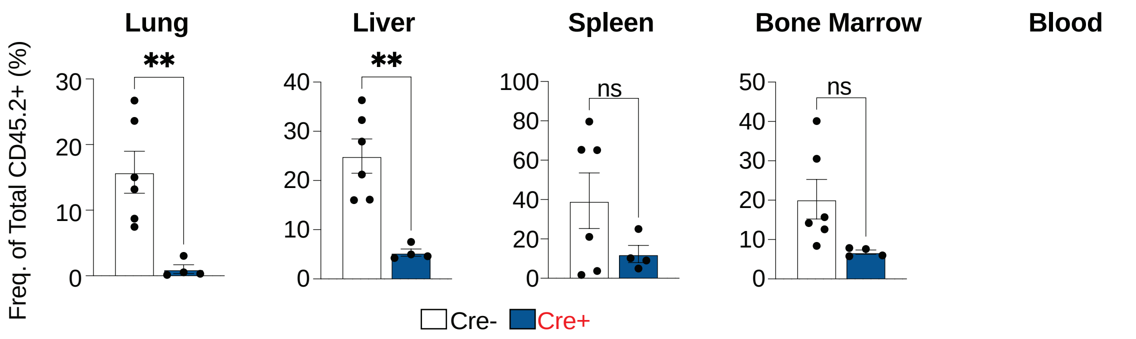


**B** Analysis of neutrophil frequencies in irradiated mice

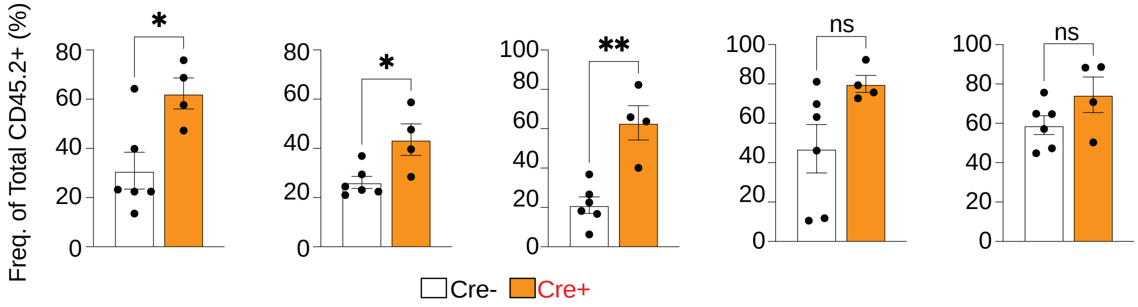


**C** Analysis of Ly6C+ monocyte frequencies in irradiated mice

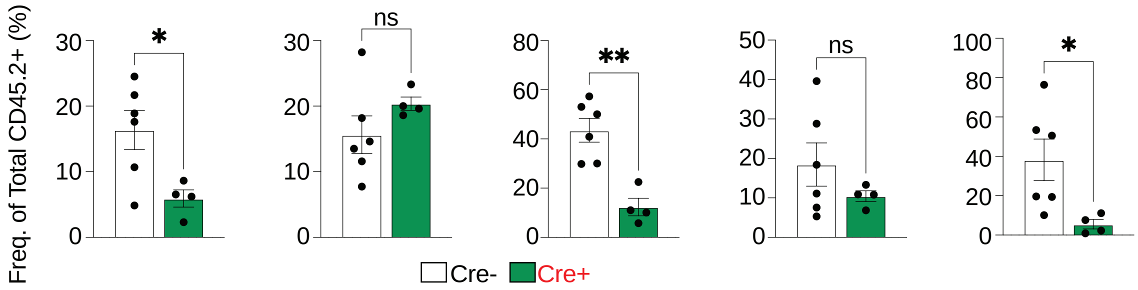


**D** Analysis of $CX_3CR1$+ monocyte frequencies in irradiated mice

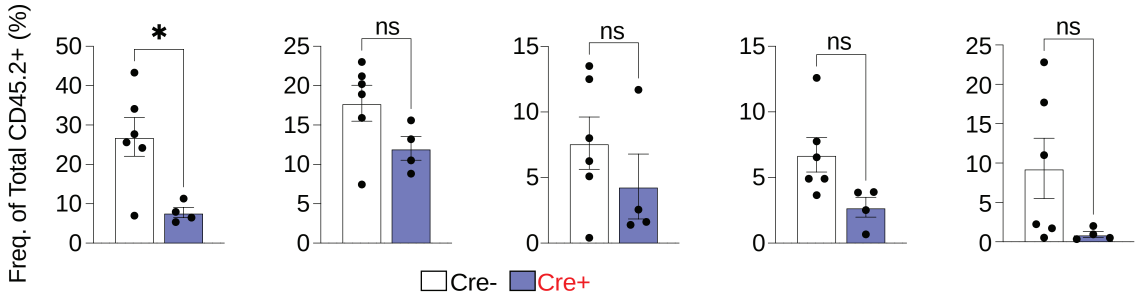

**Figure S8. Human cells do not differentiate into macrophages in the presence of WNK463.**

**(A, B)** Representative histograms and summary plots for flow cytometry analysis of human cell surface markers in human monocytes **(A)** and myeloid progenitors **(B)** treated with human M-CSF in the presence of vehicle or WNK463 as detailed in **Figure 3**. Data are from four independent experiments with two-to-three unique donors per experiment. Data are shown as mean ± SEM. Statistical significance was determined via independent samples *t*-test. *$p < .05$, **$p < .01$, ns = not significant.

**Figure S8: Human cells do not differentiate into macrophages in the presence of WNK463**

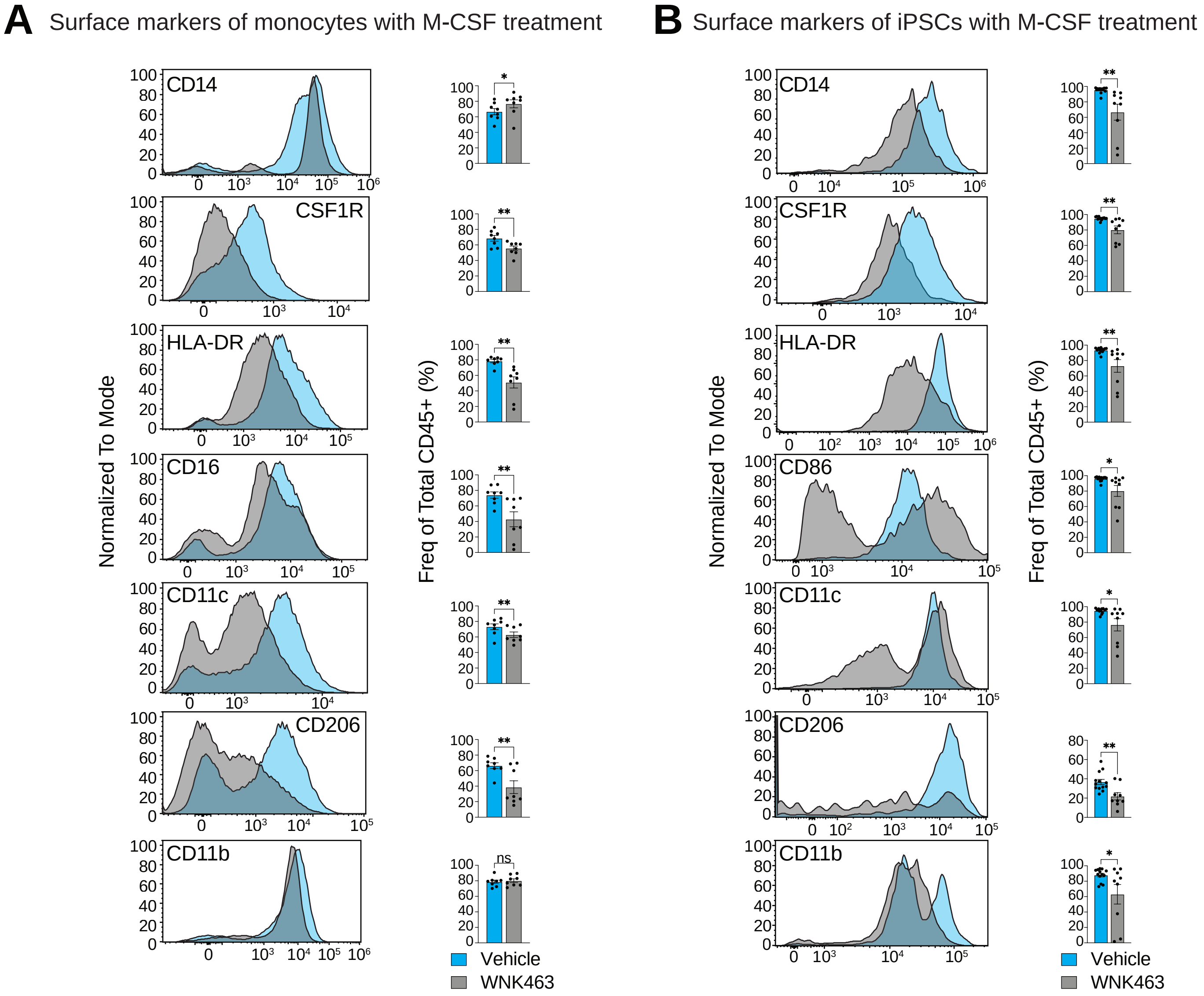
